# A novel copper-sensing two-component system for activating Dsb genes in bacteria

**DOI:** 10.1101/2020.08.17.255257

**Authors:** Liang Yu, Qiao Cao, Weizhong Chen, Nana Yang, Cai-Guang Yang, Quanjiang Ji, Min Wu, Taeok Bae, Lefu Lan

**Affiliations:** University of Chinese Academy of Sciences, No.19A Yuquan Road, Beijing, 100049, China; State Key Laboratory of Drug Research, Shanghai Institute of Materia Medica, Chinese Academy of Sciences, Shanghai 201203, China; College of Life Science, Northwest University, Xi’an, China; School of Physical Science and Technology, ShanghaiTech University, Shanghai 201210, China; Department of Biomedical Sciences, University of North Dakota, Grand Forks, North Dakota 58203-9037, USA; Department of Microbiology and Immunology, Indiana University School of Medicine-Northwest, Gary, Indiana 46408, USA; NMPA Key Laboratory for Testing Technology of Pharmaceutical Microbiology, Shanghai Institute for Food and Drug Control, Shanghai, China

**Keywords:** *Pseudomonas aeruginosa*, two-component system, disulfide bond formation, gene regulation, copper resistance

## Abstract

Copper is an essential element for biological systems but becomes toxic when present in excess. In *Pseudomonas aeruginosa*, an important human pathogen, the resistance to copper requires the induction of *dsbDEG* operon encoding proteins involved in disulfide-bond formation (Dsb). However, it is unknown how the copper stress induces the transcription of the operon. Here, we report that the exogenous copper induces the transcription of the *dsbDEG* operon through a new copper-sensing two-component system named DsbRS. The *dsbRS* is divergently transcribed from the *dsbDEG* operon, and the response regulator DsbR binds to the intergenic region between the operons. In the absence of copper, the sensor kinase DsbS acts as a phosphatase toward DsbR and thus blocks the transcription of the operons. However, in the presence of copper, the metal ion directly binds to the sensor domain of DsbS, for which the Cys82 residue plays a critical role. The copper-binding appears to inhibit the phosphatase activity of DsbS, leading to activation of DsbR. The copper resistance of the *dsbRS* knock-out mutant was restored by ectopic expression of the *dsbDEG* operon, confirming the critical role of the operon in the resistance to copper. Strikingly, cognates of *dsbRS*-*dsbDEG* pair are widely distributed across eubacteria. Also, a DsbR-binding site, which contains the consensus sequence 5’-TAA-N_7_-TTAAT-3’, is detected in the promoter region of *dsbDEG* homologs in those species. Thus, regulation of Dsb genes by DsbRS represents a novel mechanism by which bacterial cells cope with copper stress.

**Importance:** Copper is an essential redox active cofactor that becomes highly cytotoxic when present in excess. Therefore, in order to evade copper toxicity, bacteria must perceive copper stress and tightly regulate genes expression. In the present study, we identify a new copper-sensing two-component system (designated DsbRS) in *Pseudomonas aeruginosa*, an important human pathogen. We provide multiple lines of evidence that upon copper binding to the periplasmic domain of DsbS, its phosphatase activity is blocked, and the phosphorylated DsbR directly activates the transcription of a number of copper-induced genes including those involved in protein disulfide-bond formation (Dsb). This study suggests that regulation of Dsb genes by DsbRS may be an underappreciated regulatory mechanism by which bacteria sense and respond to copper.

## Introduction

Copper (Cu) is an essential micronutrient for almost all organisms. Due to its ability to cycle between two oxidation states, Cu^+^ and Cu^2+^, it is widely used in bacteria as a cofactor in a wide variety of enzymes catalyzing redox reactions (1, 2). However, the redox property of copper also creates a potential hazard to bacteria (1–3). In excess, copper is extremely toxic owing in part to its ability to generate reactive oxygen species, and there is growing interest in utilizing copper for infection control due to its antibacterial properties (1–4). Interestingly, recent studies suggest that the innate immune system of the animal host utilizes the toxic properties of copper to kill bacteria (2, 5, 6).

To evade copper toxicity, bacteria must tightly regulate genes involved in the copper homeostasis (7, 8), gathering information through various cytoplasmic Cu-sensing one-component systems (i.e., CueR, CsoR, and CopY) and periplasmic Cu-sensing two-component systems (TCSs) (i.e., CusRS, CopRS, PcoRS, and CinRS) (7–9). Among them, TCSs are the most common mechanism of transmembrane signal transduction in bacteria, and the canonical TCS system consists of a sensor histidine kinase (HK) and a cognate response regulator (RR) (10). Upon sensing a stimulus, the HK autophosphorylates at a conserved histidine and subsequently transfers this phosphoryl group to a conserved aspartate residue located in the regulatory receiver domain of an RR that coordinates changes in bacterial behavior, often through its activity as a transcriptional regulator (10, 11). Many HK proteins have both kinase and phosphatase activities, allowing them to phosphorylate and remove phosphate from the cognate RR (12, 13).

In the archetypical *Escherichia coli* TCS CusRS, Cu^+^ binding to the periplasmic loop of HK CusS promotes its autophosphorylation, and the subsequent phosphorylation of its cognate RR CusR, which then activate the transcription of *cusCFBA* genes encoding proteins for the Cu^+^ efflux pump CusCFBA to ameliorate copper toxic effects (14–19). In addition to CusRS, various periplasmic copper-sensing TCSs (e.g., CopRS, PcoRS, and CinRS) have been identified (8, 20–23), which supports the notion that the bacterial cell envelope is an important target for copper stress (7, 24). Indeed, copper can inhibit lipoprotein maturation and causes periplasmic protein misfolding due to the formation of non-native disulfide bonds (7, 24, 25). In order to evade copper toxicity, therefore, bacteria not only control intracellular copper homeostasis but also must minimize and repair copper-induced damage (7).

The bacterial cell envelope is the first line of defense against environmental threats, and the periplasmic protein maturation often involves covalent linking of cysteine residues through disulfide bonds (26). To date, the process of disulfide bond formation (Dsb) in the cell envelope of *E. coli* by the Dsb system is quite well understood (27, 28). In brief, DsbA protein transfers its disulfide bond to newly synthesized proteins and is reoxidized back to its active oxidized state by DsbB, while DsbC rearranges the misfolded proteins and is maintained in its active state form by DsbD (27, 28). Although there is an unexpected diversity in the mechanisms of disulfide bond formation (29, 30), the Dsb system appears to have critical roles in copper resistance in various bacterial species (24, 25, 31–35), including *Pseudomonas aeruginosa* (36), a notorious human pathogen that causes more than 10% infections in most hospitals (37) and is among the microorganisms for which there is an urgent need for novel antibiotics (38).

It has been reported that in *P. aeruginosa*, genes involved or potentially involved in protein disulfide bond formation and/or sulfenic acid reduction (i.e., *dsbD*, *dsbE*, and *dsbG*, forming a putative *dsbDEG* operon) (39, 40) were significantly induced by Cu (> 15-fold, in Cu-adapted population) (36). Importantly, in all cases, transposon insertions in those genes (i.e., *dsbD*, *dsbE*, and *dsbG*) decrease the resistance of *P. aeruginosa* to copper (36). Intrigued by these findings, we sought to examine how those Dsb genes are regulated by copper. In this study, we showed that the *dsbDEG* operon is a major target of a novel copper-sensing TCS (i.e., PA2479-PA2480), where PA2479 (designated <u>Dsb</u>-associated <u>R</u>egulator, DsbR) is an RR and PA2480 (designated <u>Dsb</u>-associated <u>S</u>ensor, DsbS) is a HK. We further showed that regulation of Dsb genes by DsbRS is a potentially widespread mechanism that controls the response of bacteria to copper stress.

## Results

### Copper induces the expression of *dsbDEG* and *dsbRS* operons

Using reverse transcription (RT)-PCR analysis, we showed that *dsbD*, *dsbE*, and *dsbG* do indeed constitute a transcriptional unit (Fig. S1). In order to further examine the transcriptional activation of *dsbDGE* operon by copper, we constructed a *dsbD* promoter-*lux* fusion (*dsbD*-*lux*, Table S1) and measured its activity in a wild-type (WT) *P. aeruginosa* MPAO1 strain throughout the growth period. 0.5 mM Cu^2+^ were enough to activate the *dsbD* promoter, and 1.5 mM Cu^2+^ induced a 30-fold increase in the activity of *dsbD*-*lux* in stationary growth phase (at 15 h) (Fig. 1A), indicating that the Cu-induced increased steady-state mRNA levels of *dsbDGE* (36) are likely resulted from increased transcription. In addition, we observed that *dsbD-lux* expression positively correlates with the copper toxicity effect of the growth of *P. aeruginosa* (Fig. 1A).

**Fig. 1.**
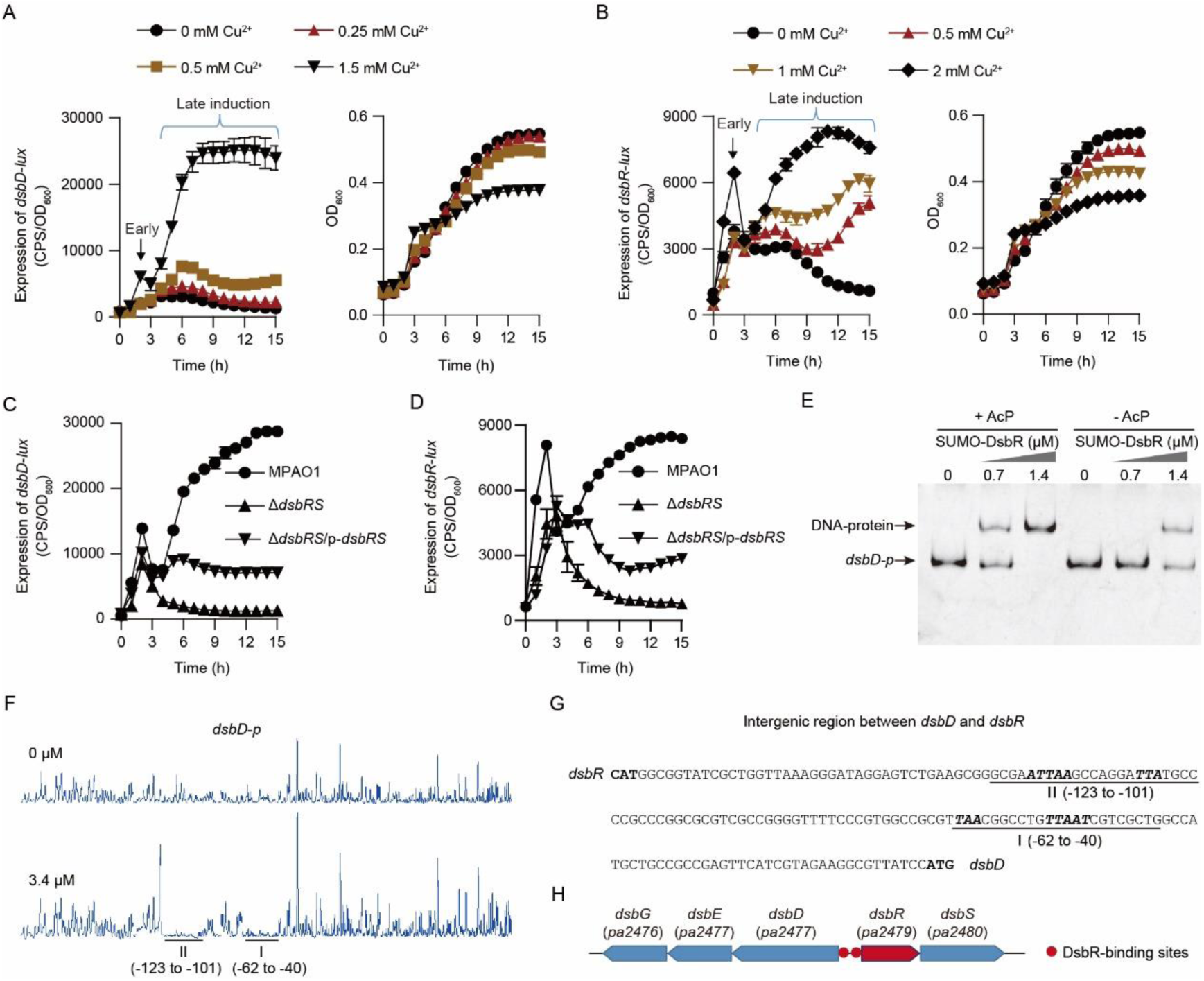
Copper induces the transcription of *dsbDEG* and *dsbRS* operons through DsbRS. (A and B) Expression of *dsbD-lux* (A) and *dsbR-lux* (B) and the corresponding growth curve (left panel) of WT MPAO1 strain cultured in LB medium supplemented without or with indicated concentrations of CuSO4. Arrow and bracket indicate the early and late Cu-induced activation of either *dsbD-lux* (A) or *dsbR-lux* (B), as indicated. The value of CPS/OD_600_ (counts per second/an optical density at 600 nm) became an indicator of the promoter activity. Data from n = 3 biological replicates reported as mean ± SD. (C and D) Expression of *dsbD*-*lux* (C) and *dsbR*-*lux* (D) in *P. aeruginosa* strains cultured in LB medium supplemented with 2 mM CuSO_4_. Wild-type (WT) MPAO1 and Δ*dsbRS* mutant strain harbor an empty pAK1900 plasmid as control; p-*dsbRS* denotes the pAK1900-*dsbRS* plasmid (Table S1). Data from n = 3 biological replicates reported as mean ± SD. (E) EMSAs show the binding of SUMO-DsbR to a 578-bp DNA fragment covering the intergenic region of *dsbD*-*dsbR* (*dsbD*-*p*) in the presence (+) or absence (-) of 30 mM AcP. (F) Dye primer-based DNase I footprint assays show the protected pattern of *dsbD*-*dsbR* intergenic region DNA after digestion with DNase I following incubation without or with purified SUMO-DsbR protein (3.4 µM). Two DsbR-protected regions are underlined (relative to the start codon of *dsbD*). (G) *dsbD*-*dsbR* intergenic region DNA sequence with a summary of the DNase I footprint assay results. The consensus sequence (i.e., TAA-N_7_-TTAAT) shared by the two DsbR-protected regions are in bold and in italics. The start codon of *dsbR* and *dsbD* are in bold. (H) Schematic representation of *dsbDEG* operon, *dsbRS* operon, and DsbR-binding sites.

On the *P. aeruginosa* genome, the *dsbDEG* operon is organized upstream and is transcribed divergently from the *pa2479*-*pa2480* operon (hereinafter referred to *dsbRS*) (Fig. S1A, also see in Fig. 1H) that encodes a TCS with uncharacterized biological and biochemical functions (www.pseudomonas.com). These two operons were often co-regulated (36, 41–44), implying that copper may also have a role in the transcriptional activation of *dsbRS*. Using promoter fusion experiments, we were able to demonstrate that copper increased the promoter activity of *dsbRS* (*dsbR-lux*) (Fig. 1B). As shown, when exposure to 2 mM Cu^2+^, expression levels of the *dsbR-lux* increased rapidly and reached a peak at 2 h with a 1.7-fold induction (hereinafter designated as early induction), and then declined by approximately 50 % over a 1-h period (Fig. 1B). After that, the expression level of *dsbR-lux* gradually increased again (hereinafter designated as late induction) and reached a 7.5-fold induction by Cu^2+^ after 12 h (Fig. 1B). A similar expression pattern was also observed for the Cu-induced *dsbD-lux* (Fig. 1A).

### DsbR binds between the *dsbDEG* and the *dsbRS* operon

The co-regulation of *dsbDGE* and *dsbRS* (36, 41–44) also suggests a potential regulative role of DsbRS for *dsbDGE*. To test this hypothesis, we generated a *dsbRS* deletion mutant (Δ*dsbRS*) and measured the *dsbD-lux* activity in a WT MPAO1, a Δ*dsbRS*, and a Δ*dsbRS*-complemented strain (Δ*dsbRS*/p-*dsbRS*). Deletion of *dsbRS* severely abolished the *dsbD*-*lux* activity in *P. aeruginosa* MPAO1 strain grown in LB medium supplemented with 2 mM CuSO_4_ (Fig. 1C). The introduction of a plasmid carrying *dsbRS* (p-*dsbRS*) into the *dsbRS* deletion mutant (Δ*dsbRS*) partially restored the wild-type characteristics of *dsbD-lux* activity, indicating that DsbRS activates the transcription of *dsbDEG*. A similar result was obtained when the promoter activity of *dsbR* (*dsbR-lux*) was examined (Fig. 1D).

Like many RRs (45, 46), DsbR can be phosphorylated by acetyl phosphate (AcP) (Fig. S2 A and B). Using electrophoretic mobility shift assays (EMSAs), we showed that the purified recombinant DsbR protein (i.e., SUMO-DsbR) is phosphorylated (Fig. S2 C and D) and could shift a DNA fragment (*dsbD-p*) covering the intergenic region between *dsbD* and *dsbR* (Fig. S2E), although it failed to do this for an unrelated DNA fragment that serves as a negative control (Fig. S2F). Acetyl phosphate also can enhance the DsbR binding to its target *dsbD-p* (Fig. 1E), indicating that DsbR is activated by phosphorylation. Using a dye-primer-based DNase I footprint assay, we uncovered two DsbR-protected regions in the intergenic region between *dsbD* and *dsbR* (Fig. 1F). Interestingly, all the two DsbR-protected regions contain a consensus sequence TAA-N_7_-TTAAT (where N refers to any nucleotide) (Fig. 1G). Based on these results, we concluded that phosphorylated DsbR binds to and activate the promoter of *dsbDEG* operon and its own promoter (Fig. 1H).

### *dsbR* activates the expression of *dsbDEG* and *dsbRS*, whereas *dsbS* inhibits it

To further investigate the roles of DsbRS in the promoter activity of *dsbDEG*, we measured the activity of *dsbD*-*lux* in a WT MPAO1 strain, a Δ*dsbRS* mutant strain, and a Δ*dsbRS* mutant strain carrying either p-*dsbR*, p-*dsbS* or p-*dsbRS* plasmid (Table S1). When bacteria were grown in LB medium without external addition of copper, the activity of *dsbD-lux* in the Δ*dsbRS* mutant was low and comparable to those in the WT MPAO1 strain (Fig. 2A); however, the introduction of p-*dsbR*, but not p-*dsbS* or p-*dsbRS*, into the Δ*dsbRS* mutant increased the activity of *dsbD*-*lux* (>100-fold). A similar phenomenon was observed for the *dsbR-lux* transcriptional reporter fusion (Fig. 2A). These observations indicate that DsbR is functional in the absence of DsbS, also suggesting that DsbS inhibits DsbR under the testing conditions. Further supporting these notions, deletion of *dsbS* alone caused over-expression of either *dsbD*-*lux* (a 71-fold increase) or *dsbR*-*lux* (a 6-fold increase) (Fig. 2A).

**Fig. 2.**
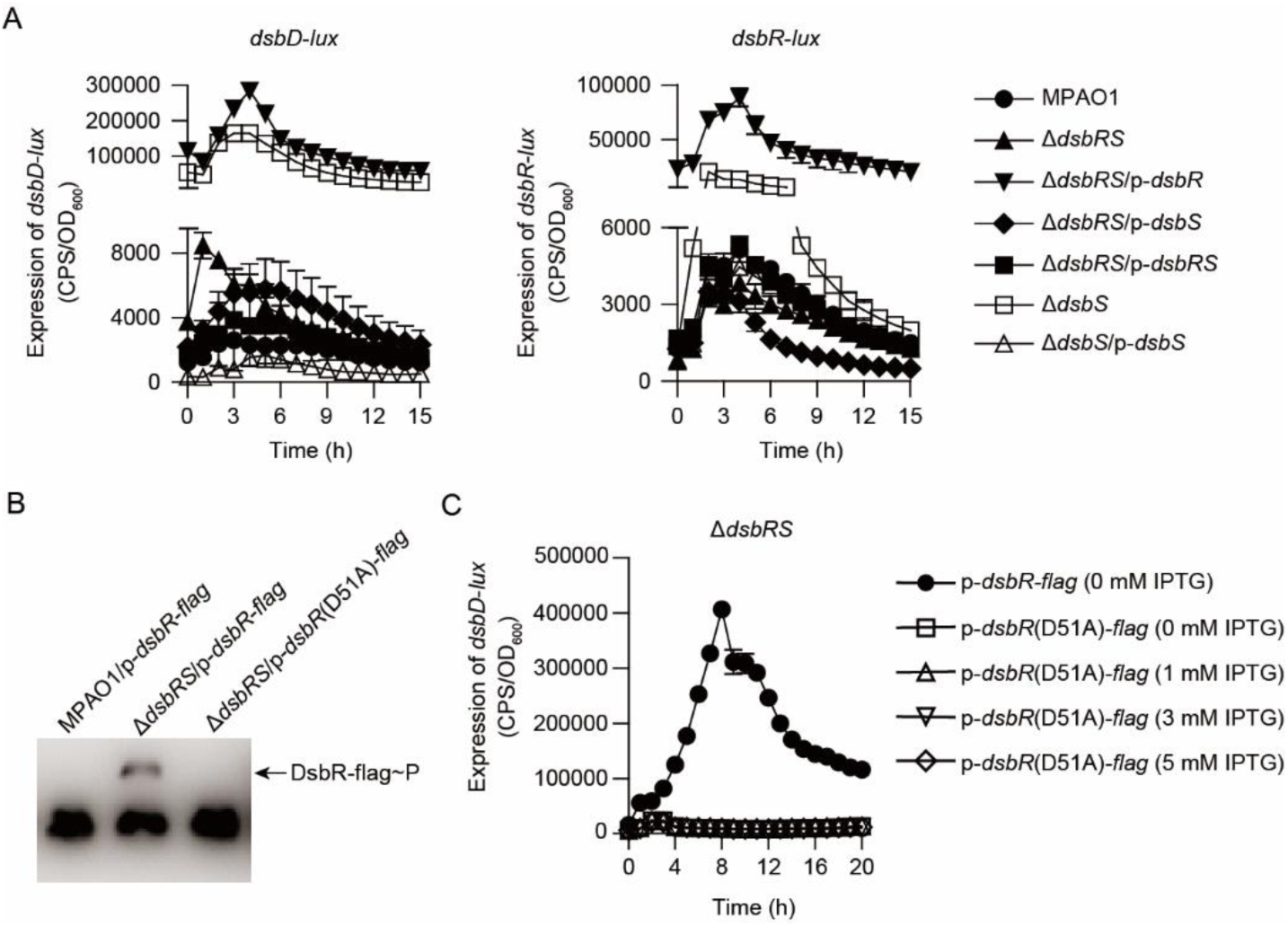
DsbS inhibits DsbR in the absence of copper stimulus. (A) Expression of *dsbD*-*lux* (left panel) and *dsbR*-*lux* (right panel) in *P. aeruginosa* strains grown in LB medium. MPAO1, Δ*dsbS*, and Δ*dsbRS* harbor an empty pAK1900 plasmid as control; p-*dsbR*, p-*dsbS*, and p-*dsbRS* denote pAK1900-*dsbR*, pAK1900-*dsbS*, and pAK1900-*dsbRS* (Table S1), respectively. Data from n = 3 biological replicates reported as mean ± SD. (B) Phos-tag analysis of the phosphorylation level of DsbR-flag and DsbR(D51A)- flag. Protein samples were derived from bacteria grown in LB at 37°C for 5 h. (C) Effect of D51A missense mutation in *dsbR* on its ability to promote the expression of *dsbD*-*lux*. The expression of *dsbD*-*lux* was examined in *P. aeruginosa* strains grown in LB supplemented without or with indicated concentrations of isopropyl-1-thio-β-d-galactopyranoside (IPTG). Data from n = 3 biological replicates reported as mean ± SD.

### The phosphorylation at D51 seems to be essential for DsbR to activate transcription of target genes

Since *dsbS* deficiency caused an increase in the regulatory activity of DsbR (Fig. 2A), we sought to determine if the phosphorylation status of DsbR is affected by the absence of *dsbS*, given that DsbR appears to be activated by phosphorylation *in vitro* (Fig. 1E, Fig. S2). With a plasmid (p-*dsbR-flag*, Table S1), Flag-tagged DsbR proteins were expressed in WT MPAO1 strain and the Δ*dsbRS* mutant; then, the total bacterial proteins were resolved on SDS-PAGE gels containing Phos-tag^TM^ acrylamide and subjected to Western blotting with an anti-FLAG antibody. In the WT strain, only the non-phosphorylated DsbR-flag was observed, whereas, in the Δ*dsbRS* mutant, phosphorylated DsbR-flag (DsbR-flag∼P) was also detected (Fig. 2B). These results suggest that the phosphorylation level of DsbR is increased upon the absence of DsbS. Additionally, when the predicted phosphorylation site Asp51 was substituted with alanine, the *in vivo* phosphorylation of DsbR was eliminated (Fig. 2B), supporting the idea that DsbR is phosphorylated at Asp51. Moreover, although DsbR-flag was able to largely increase the expression of *dsbD-lux* in a Δ*dsbRS* mutant strain (Fig. 2C), the DsbR^D51A^-flag (Asp51 replaced by alanine) failed to do this (Fig. 2C) regardless of its protein levels (Fig. S3A), indicating that phosphorylation of DsbR at Asp51 is essential for the activation of DsbR in *P. aeruginosa*. Thus, DsbS appears to inhibit DsbR in the absence of copper stimulus.

### S239A mutation increases the autokinase activity of DsbS^CTD^ but decrease its phosphatase activity against DsbR

DsbS is a member of the HisKA subfamily of bacterial membrane-associated histidine kinases (Fig. 3A), and the His235 and Ser239 residues are predicted to be required for its autokinase and phosphatase activities, respectively (47). Indeed, in an autophosphorylation assay with purified truncated DsbS protein (i.e., DsbS^CTD^, the C-terminal region of DsbS, residues 225-440, where the insoluble N-terminal membrane spanning domains is replaced by a His_6_ Ni-affinity tag), the substitution of the His235 with alanine (H235A) completely abolished the autophosphorylation (Fig. S3B), confirming that His235 is the conserved autophosphorylation site at DsbS. In contrast to the H235A substituent, replacement of the Ser239 with alanine (S239A) increased the autokinase activity of DsbS^CTD^ (Fig. 3B).

**Fig. 3.**
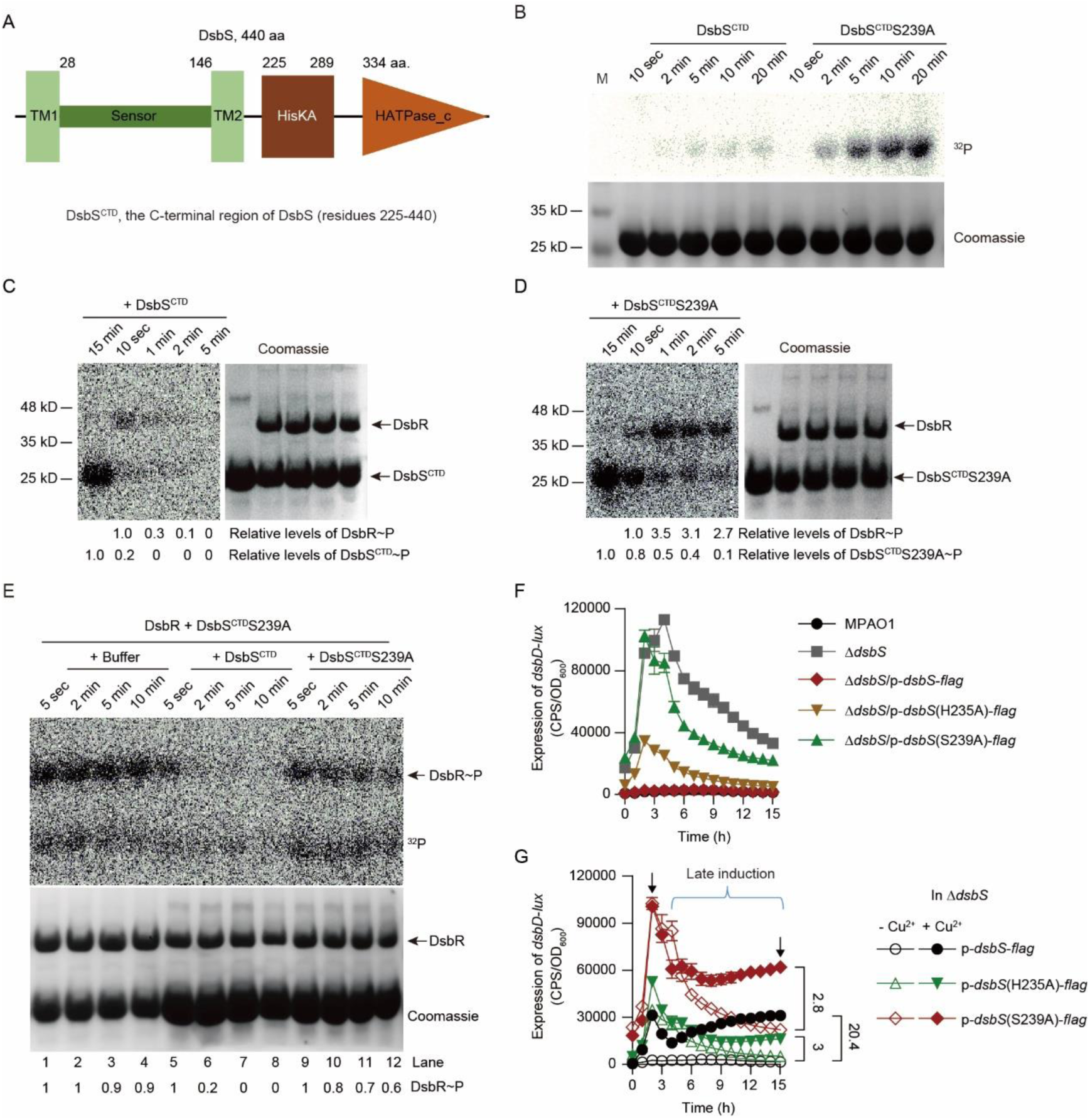
DsbS acts as a bifunctional histidine sensor kinase. (A) Schematic view of the secondary structure of DsbS predicted by SMART (http://smart.embl-heidelberg.de/). TM indicates transmembrane helix. (B) Time course of auto-phosphorylation of N-terminal His_6_-tagged DsbS cytoplasmic domain (i.e., DsbS^CTD^) and its S239A (i.e., DsbS^CTD^S239A) mutant incubated with [γ-^32^P]ATP. M, protein marker. Samples were fractionated by 12% SDS-PAGE (lower panel), and radiolabeled proteins were visualized by autoradiography (upper panel). (C and D) Transphosphorylation of SUMO-DsbR by DsbS^CTD^∼P (C) and DsbS^CTD^S239A∼P (D). DsbS^CTD^ or DsbS^CTD^S239A was first auto-phosphorylated for 15 min in the presence of [γ-^32^P]ATP (lane 1), and then purified SUMO-DsbR proteins were added to the reaction assay and incubated at room temperature for the indicated time. For quantifications, the signal intensity of phosphorylation (left panel) was normalized to the intensity obtained with Coomassie Blue stained gel (used as loading control, right panel) and the results are reported as fold changes with control sample set to 1, as indicated. (E) Time course of de-phosphorylation of SUMO-DsbR∼P. SUMO-DsbR was first transphosphorylated by DsbS^CTD^S239A for 15 min. Control solutions (+ Buffer, lane 1-4), DsbS^CTD^ (+ DsbS^CTD^, lane 5-8), or DsbS^CTD^S239A (+ DsbS^CTD^S239A, lane 9-12) was added to the reaction mixtures and samples were taken at the indicated time points and analyzed by SDS-PAGE. The results are reported as fold changes with phosphorylation level of DsbR at 5 sec in each experimental group (lane 1, 5, and 9) set to 1, respectively. (F) Expression of *dsbD*-*lux* in *P. aeruginosa* strains grown in LB medium. MPAO1 and Δ*dsbS* harbor an empty pAK1900 plasmid as control; p-*dsbS*-*flag*, p-*dsbS*^H235A^-*flag*, and p-*dsbS*^S239A^-*flag* denote pAK1900-*dsbS*-*flag*, pAK1900-*dsbS*^H235A^-*flag*, and pAK1900-*dsbS*^S239A^-*flag*, respectively (Table S1). Data from n = 3 biological replicates reported as mean ± SD. (G) Expression of *dsbD*-*lux* in Δ*dsbS* mutant carrying indicated *dsbS* variants. Bacteria were grown in LB supplemented without (- Cu^2+^) or with 2.5 mM CuSO_4_ (+ Cu^2+^). Left arrow and bracket respectively indicate the early and late activation of *dsbD*-*lux* by copper. Right arrow indicates the late activation of *dsbD*-*lux* at 15 h. Data from n = 3 biological replicates reported as mean ± SD.

Because DsbS and DsbR were predicted to form a TCS (www.pseudomonas.com), we sought to assess the ability of DsbS to undergo phosphotransfer to DsbR. For this purpose, DsbS^CTD^ was autophosphorylated with [γ- ^32^P]ATP for 15 min, and then purified SUMO-DsbR was added to the reaction mixture and sampled at fixed sampling intervals thereafter. Although SUMO-DsbR was not phosphorylated by [γ-^32^P]ATP (Fig. S3C), it was trans-phosphorylated by DsbS^CTD^∼P (Fig. 3C). Interestingly, DsbR phosphorylation waned over times rapidly and it was undetectable at five minutes (Fig. 3C), indicating that DsbS not only acts as a kinase but also a phosphatase against DsbR. In contrast, although DsbS^CTD^S239A∼P was able to phosphorylate DsbR, it failed to dephosphorylate phosphorylated DsbR (DsbR∼P) over a five-minute time course (Fig. 3D), indicating that replacement of Ser239 with alanine (S239A) diminishes the phosphatase activity of DsbS.

When SUMO-DsbR was transphosphorylated by DsbS^CTD^S239A∼P, the SUMO-DsbR∼P showed a half-life more than 10 min (Fig. 3E, lane 1-4). However, when DsbS^CTD^ was added to the reaction mixture in a 2:1 (DsbS^CTD^/SUMO-DsbR) molar ratio, the half-life of SUMO-DsbR∼P decreased to less than 2 min (Fig. 3E, lane 5-8), further supporting that DsbS has significant phosphatase activity towards DsbR∼P. In contrast, when DsbS^CTD^S239A was added in a 2:1 (DsbS^CTD^S239A /SUMO-DsbR) molar ratio, the half-life of SUMO-DsbR∼P was more than 10 min (Fig. 3E, lane 9-12). These results, together with the *in vivo* data (Fig. 2), suggest that DsbS is bifunctional, acting as a phosphatase toward phosphorylated DsbR in the absence of copper stimulus.

### His235 and Ser239 of DsbS are crucial for Cu-induction of *dsbD-lux*

As described above, His235 is required for the autokinase activity while Ser239 is critical for the phosphatase activity of DsbS^CTD^ (Fig. S3B, Fig. 3D and E). In order to investigate the autokinase and phosphatase activities of DsbS on the activation of DsbR, we examined the role of His235 and Ser239 residues in the expression of *dsbD-lux*. The plasmids containing genes encoding Flag-tagged WT DsbS or the mutant proteins DsbS(H235A) and DsbS(S239A) were introduced into a Δ*dsbS* mutant; then, the expression of *dsbD-lux*, an indicator for the regulatory activity of DsbR (Fig. 2A), was measured. When bacteria were grown in LB medium, a Flag-tagged *dsbS* gene (p-*dsb-flag*, Table S1) was able to decrease the *dsbD-lux* expression in the Δ*dsbS* mutant to a WT level (Fig. 3F), indicating that the Flag-tagged DsbS is functional and can inhibit the activation of DsbR as excepted. However, although H235A and S239A substituents had no obvious effect on the production of DsbS-Flag (Fig. S3D), they reduced the ability of Flag-tagged DsbS to inhibit the expression of *dsbD-lux* by approximately 31% and 91% respectively (determined by the maximal level of *dsbD*-*lux* activity, Fig. 3F). These results suggest that the phosphatase activity of DsbS is essential for its inhibitory effect on DsbR. In addition, the decreased ability of DsbS(H235A) to inhibit DsbR may also result from reduced phosphatase activity, which is similar to EnvZ from *E. coli* where the conserved histidine is not required but enhances phosphatase activity (48).

We observed that like WT MPAO1 strain, the Δ*dsbS*/p-*dsbS-flag* strain (i.e., Δ*dsbS* mutant carrying p-*dsbS-flag*) exhibits very low activity of *dsbD*-*lux* in the absence of external addition of copper (Fig. 3F and G). However, in the presence of Cu^2+^ (2.5 mM), the expression levels of *dsbD-lux* increased rapidly and reached a peak at 2 h with a 11.7-fold induction, and then declined by 58 % over a 2-h period (Fig. 3G). After that, the expression level of *dsbD-lux* gradually increased again, and reached a 20.4-fold Cu-induction at 15 h (Fig. 3G). Notably, H235A substituent caused an 87.1% decrease in the early (2 h) Cu-induction and an 85.3% decrease in the late (15 h) Cu-induction of *dsbD*-*lux* (Fig. 3G), indicating that the autokinase activity of DsbS is crucial for Cu-induced activation of DsbR. To our surprise, the early Cu-induction of *dsbD*-*lux* was completely abolished upon replacement of Ser239 with alanine (S239A) (Fig. 3G), suggesting that copper must inhibit the phosphatase activity of DsbS in order to evoke an early activation of DsbR. Like H235A, S239A substituent also severely reduced the late Cu-induction of *dsbD-lux* (Fig. 3G). Together with the biochemical data (Fig. 3 B-E, Fig. S3B), these results suggest that both the autokinase and phosphatase activities of DsbS are very important for the Cu-induced activation of DsbR.

### The sensor domain is required for the Cu-mediated activation of *dsbD* transcription and directly binds to Cu and Ni

In many TCSs, signal recognition and transmission occurs via a ligand binding in the periplasmic sensor domain of the HK (11). To gain insight into the role of the periplasmic domain of DsbS, we deleted the sequence encoding the periplasmic domain (residues 28-146) of flag-tagged *dsbS*, resulting in *dsbS*^Del-SD^*-flag* (Fig. 4A). Introduction of a plasmid expressing either WT *dsbS-flag* (i.e., p-*dsbS-flag*) or *dsbS*^Del- SD^*-flag* (i.e., p-*dsbS*^Del-SD^*-flag*) into a Δ*dsbS* mutant decreased the expression of *dsbD-lux* to WT levels when bacteria were cultured in LB medium (Fig. 4B), indicating that the periplasmic domain is not essential for the repressive activity of DsbS on DsbR. Importantly, although Cu^2+^ was able to increase the expression of *dsbD-lux* in the Δ*dsbS*/p-*dsbS-flag* strain, it failed to do this for the Δ*dsbS*/p-*dsbS*^Del-SD^*-flag* strain (Fig. 4C), suggesting that DsbRS requires the periplasmic domain of DsbS to exert its regulatory effect on *dsbD-lux* in response to copper.

**Fig. 4.**
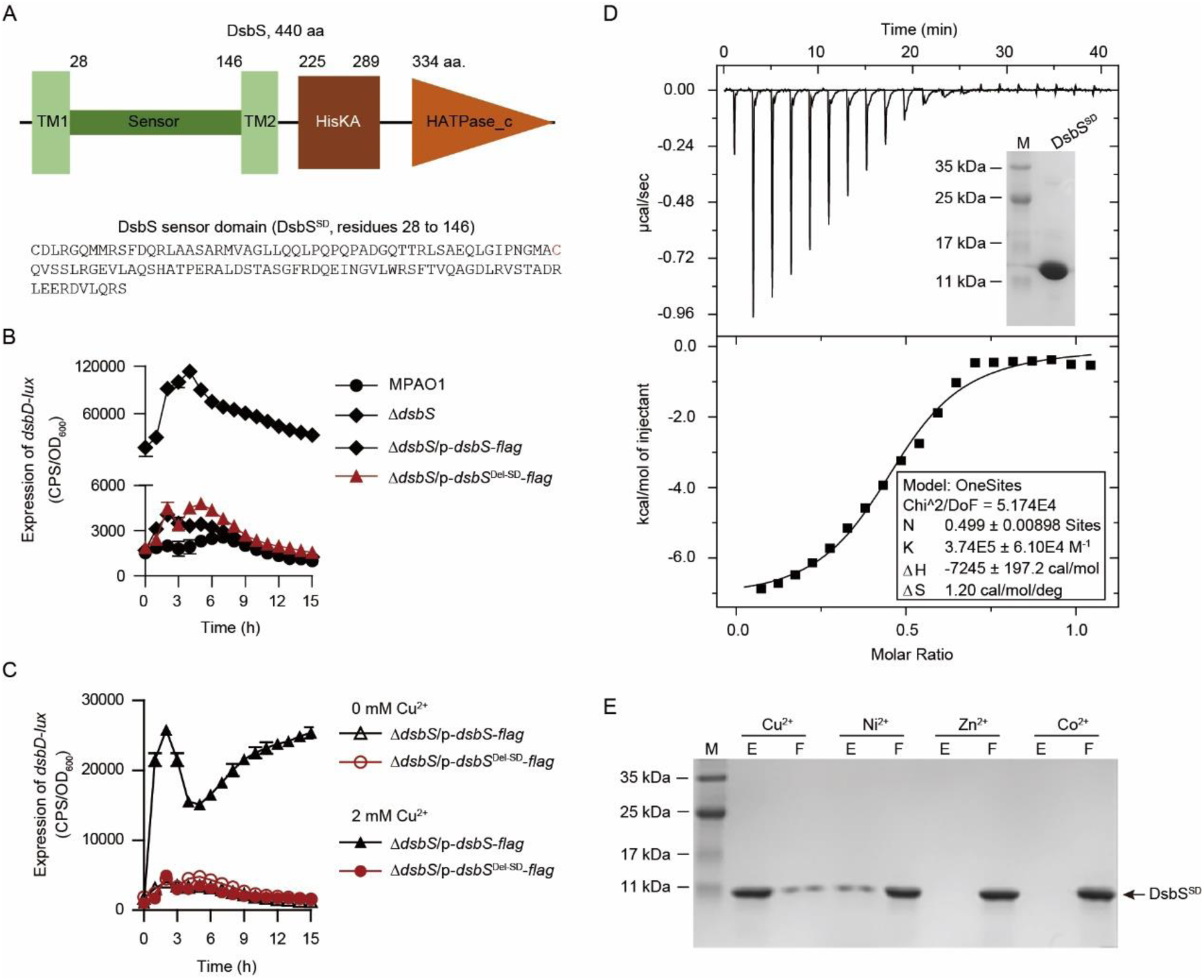
DsbS requires its periplasmic domain to sense copper. (A) Schematic view of the secondary structure of DsbS predicted by SMART (http://smart.embl-heidelberg.de/). The amino acid sequence for the N-terminal periplasmic sensor domain (DsbS^SD^, residues 28 to 146) are shown at the bottom of the panel. (B) Effect of the deletion of *dsbS*^SD^ (encoding the sensor domain of DsbS) on the expression of *dsbD-lux* in *P. aeruginosa* strains grown in LB. MPAO1 and Δ*dsbS* harbor the control plasmid pAK1900; p-*dsbS*-*flag* denotes pAK1900-*dsbS*-*flag*; p-*dsbS*^Del-SD^-*flag* denotes pAK1900-*dsbS*^Del-SD^-*flag* (Table S1). Data from n = 3 biological replicates reported as mean ± SD. (C) Effect of the deletion of *dsbS*^SD^ on the copper-induced activation of *dsbD-lux*. Data from n = 3 biological replicates reported as mean ± SD. (D) Monitoring Cu^2+^ binding to DsbS^SD^ using isothermal titration calorimetry. Top, raw data; shown in the figure insert is the image of SDS-PAGE gel of the tag-free DsbS^SD^; M, protein marker. Bottom, plot of integrated heats versus the Cu/DsbS^SD^ ratio. The curve represents the best fit for a one-site binding model. (E) Analysis of the interactions of metal ions with DsbS^SD^ using immobilized metal ion affinity chromatography (IMAC). His-Bind resin columns were loaded with 0.5 mM CuSO_4_, NiSO_4_, ZnSO_4_, or CoCl_2_, as indicated, and about 10 µg of purified DsbS^SD^ were applied to the columns. After centrifugation the flowthroughs (F) were removed and the resins were washed three times with 1 ml of buffer R in order to remove the non-bound proteins. The protein was further eluted with 100 µl of 0.4 M imidazole in buffer R, and 10 µl of flowthrough (F) or eluate (E) was loaded onto an SDS-PAGE gel.

To further examine the role of the sensor domain in the Cu-sensing by DsbS, we expressed and purified the periplasmic domain of DsbS (i.e., DsbS^SD^). In an isothermal titration calorimetry (ITC) assay, DsbS^SD^ bound to Cu^2+^ with an affinity of 2.7 µM (Fig. 4D), indicating that DsbS directly mediates the Cu-binding of DsbS. The immobilized metal ion affinity chromatography (IMAC) showed that DsbS^SD^ could bind Cu^2+^ and Ni^2+^ but not Zn^2+^ or Co^2+^ (Fig. 4E). Importantly, like Cu^2+^, Ni^2+^ was able to increase the expression of *dsbD-lux* (Fig. S4A) whereas other metal ions including Zn^2+^ and Co^2+^ were not (Fig. S4B). These data suggest that the periplasmic domain of DsbS mediates the copper-sensing *via* direct binding to copper.

### The C82 residue is critical for metal-binding of DsbS^SD^ and the response to Cu signal

In the copper binding proteins, the cysteine residue is commonly involved in copper binding (49). Indeed, DsbS^SD^ has two cysteine residues, Cys28 and Cys82 (Fig. 4A). Cys28 is adjacent to the transmembrane domain (TM1) while Cys82 locates in the central part of the periplasmic domain of DsbS. To examine the role of Cys82 in the copper-binding, we replaced Cys82 with alanine and subjected the mutant protein to the IMAC analysis. As shown, the alanine replacement of Cys82 abolishes the ability of DsbS^SD^ to bind to either Cu^2+^ or Ni^2+^ (Fig. 5A). Moreover, although the C82A substituent did not affect the production of DsbS-flag protein (Fig. S5A) or the ability of DsbS-flag to inhibit *dsbD-lux* expression in a Δ*dsbS* mutant grown in LB (Fig. 5B), it completely abolished the ability of DsbS-flag to respond to Cu (Fig. 5C). Thus, Cys82 is required for DsbS to mediate copper-induced activation of DsbR, further supporting the notion that DsbS is a copper-sensing HK (Fig. 4).

**Fig. 5.**
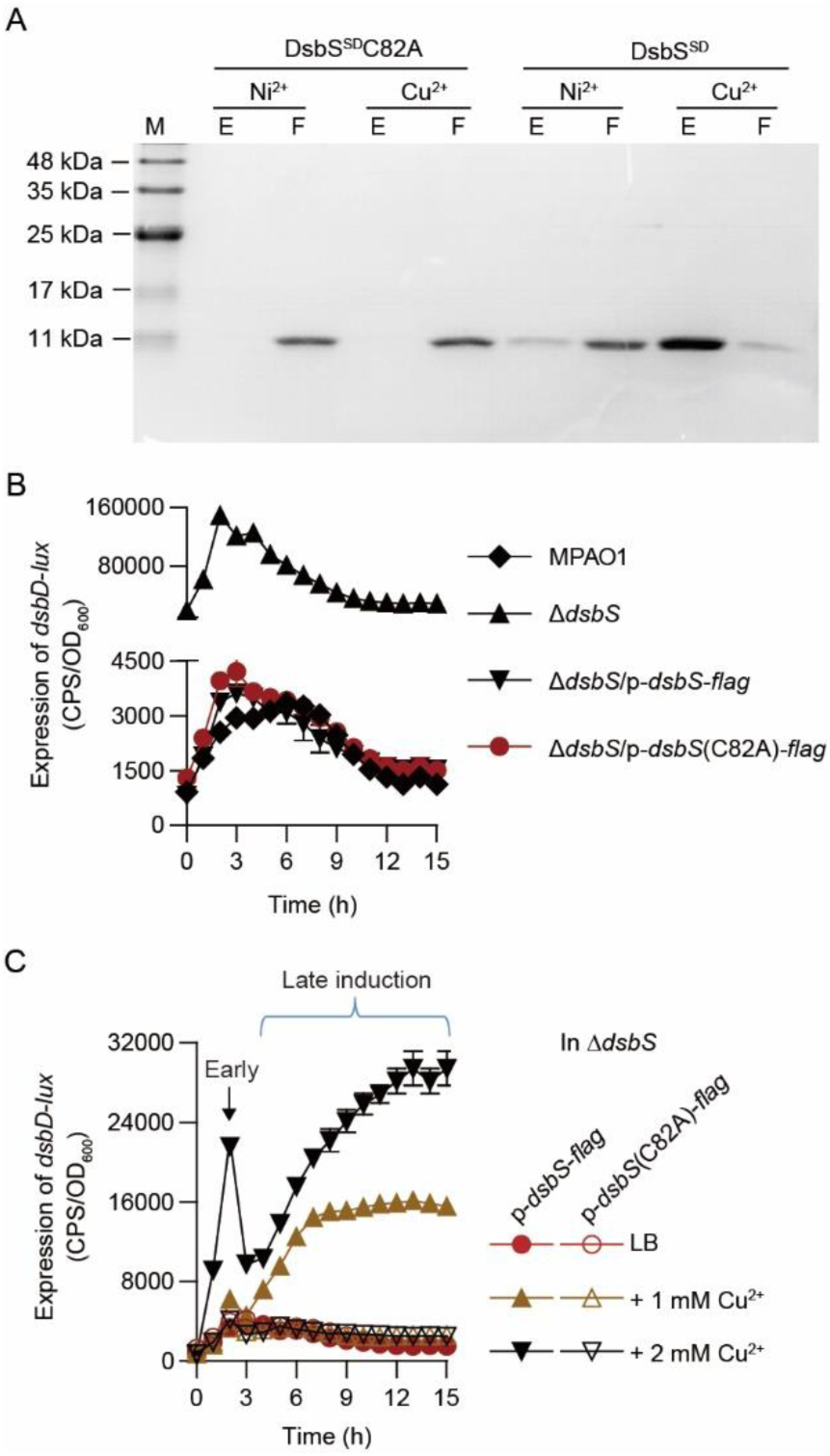
Cys82 is essential for the signaling ability of DsbS. (A) Effect of C82A substituent on the interaction of DsbS^SD^ with metal ions. His-Bind resin columns were loaded with 0.5 mM NiSO_4_ or CuSO_4_, as indicated, and about 10 µg of purified DsbS^SD^ and DsbS^SD^C82A were applied to the columns. With indicated washing and eluting, the flowthrough (F) or eluate (E) were analyzed by SDS-PAGE. (B) Effect of C82A missense mutation in *dsbS* on the expression of *dsbD-lux* in *P. aeruginosa* strains grown in LB. MPAO1 and Δ*dsbS* harbor the control plasmid pAK1900; p-*dsbS*-*flag* denotes pAK1900-*dsbS*-*flag*; p-*dsbS*(C82A)-*flag* denotes pAK1900-*dsbS*(C82A)-*flag* (Table S1). Data from n = 3 biological replicates reported as mean ± SD. (C) Effect of C82A missense mutation in *dsbS* on the copper-induced activation of *dsbD-lux*. Data from n = 3 biological replicates reported as mean ± SD.

### The *dsbRS*-mediated induction of *dsbDEG* is critical for the resistance to Cu stress

In line with the observation that DsbR can be activated in a DsbS-independent manner (Fig. 2A), the introduction of a plasmid-borne *dsbR* (p-*dsbR*) into the Δ*dsbRS* mutant increased its copper resistance phenotype (Fig. 6A), as determined by recording their growth (OD_600_) in LB broth for 15 h. Likewise, introduction of a p-*dsbRS* plasmid, but not a p-*dsbS* plasmid, was also able to increase the resistance of Δ*dsbRS* mutant towards copper (Fig. 6A), demonstrating a key role of DsbR in the copper resistance. We also found that the copper resistance phenotype of Δ*dsbRS* mutant mimics that of the Δ*dsbDEG* mutant (Fig. 6B). Interestingly, complementation of the Δ*dsbRS* mutant with a plasmid-borne *dsbDEG* operon could restore its copper resistance phenotype to WT level (Fig. 6B), suggesting that the transcriptional activation of *dsbDEG* operon by DsbRS is important for the copper resistance in *P. aeruginosa*.

**Fig. 6.**
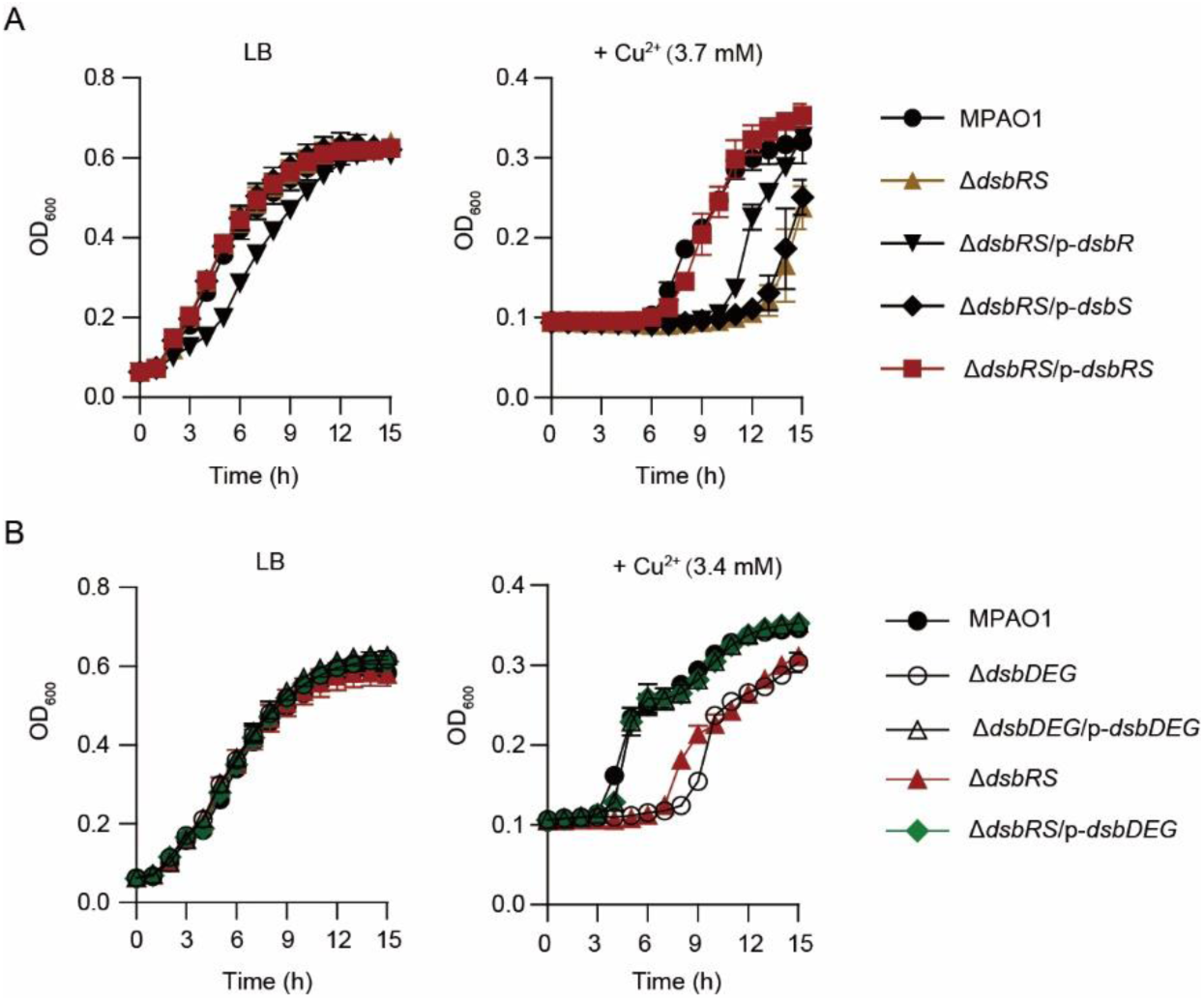
Effect of Cu^2+^ on the growth of *P. aeruginosa*. (A) Growth of *P. aeruginosa* strains were detected in LB supplement without (left panel) or with 3.7 mM CuSO_4_ (right panel). MPAO1 and Δ*dsbRS* harbor an empty pAK1900 plasmid as control; p-*dsbR*, p-*dsbS*, and p-*dsbRS* denote pAK1900-*dsbR*, pAK1900-*dsbS*, and pAK1900-*dsbRS*, respectively (Table S1). Data from n = 3 biological replicates reported as mean ± SD. (B) Growth of *P. aeruginosa* strains were detected in LB supplement without (left panel) or with 3.4 mM CuSO_4_ (right panel). MPAO1, Δ*dsbRS*, and Δ*dsbDEG* harbor the control plasmid pAK1900; p-*dsbDEG* denotes a pAK1900-*dsbDEG* plasmid (Table S1). Data from n = 3 biological replicates reported as mean ± SD.

### Genome-wide identification of DsbR targets

To obtain a more complete picture of the mechanism of action of DsbRS, we performed DsbR ChIP-seq in a Δ*dsbRS*::*dsbR-flag* strain (Δ*dsbRS* carrying an integration-proficient vector, mini-*dsbR-flag*, for the expression of *dsbR-flag*, Table S1). We identified 17 reproducible peaks of DsbR binding (Data set 1, sheet 1). Among the DsbR-bound regions, approximately 82% (14 peaks) were in the promoter regions of 22 genes (within 600 bp upstream of the translational start site) involved or potentially involved in various biological processes, including protein folding (e.g., *dsbDEG* operon and *dsbB*), protein secretion (e.g.., *hxcU* and *hxcP*, transport (e.g., *opdT*, *pa2911*, and *pstS*), and regulation of transcription (i.e., *dsbRS*, *pa3398*, *dnr*) (Data set 1, sheet 1).

Strikingly, the most enriched CHIP-seq peak was the intergenic region between *dsbD* and *dsbR* (Fig. 7A and B), suggesting that DsbR strongly binds to the promoters of *dsbDEG* and *dsbRS* in *P. aeruginosa*. Interestingly, the second-most enriched peak was the promoter region of two divergently transcribed genes including *dsbB* that plays a key role in disulfide bond formation (28, 50, 51) (Fig. 7C). In addition, ChIP-seq peaks for DsbR were also highly enriched for the promoter regions of genes (i.e, *opdT*, *pa2506*, and *pa4803*) (Fig. 7 D and E) whose expression was largely upregulated by copper (36).

**Fig. 7.**
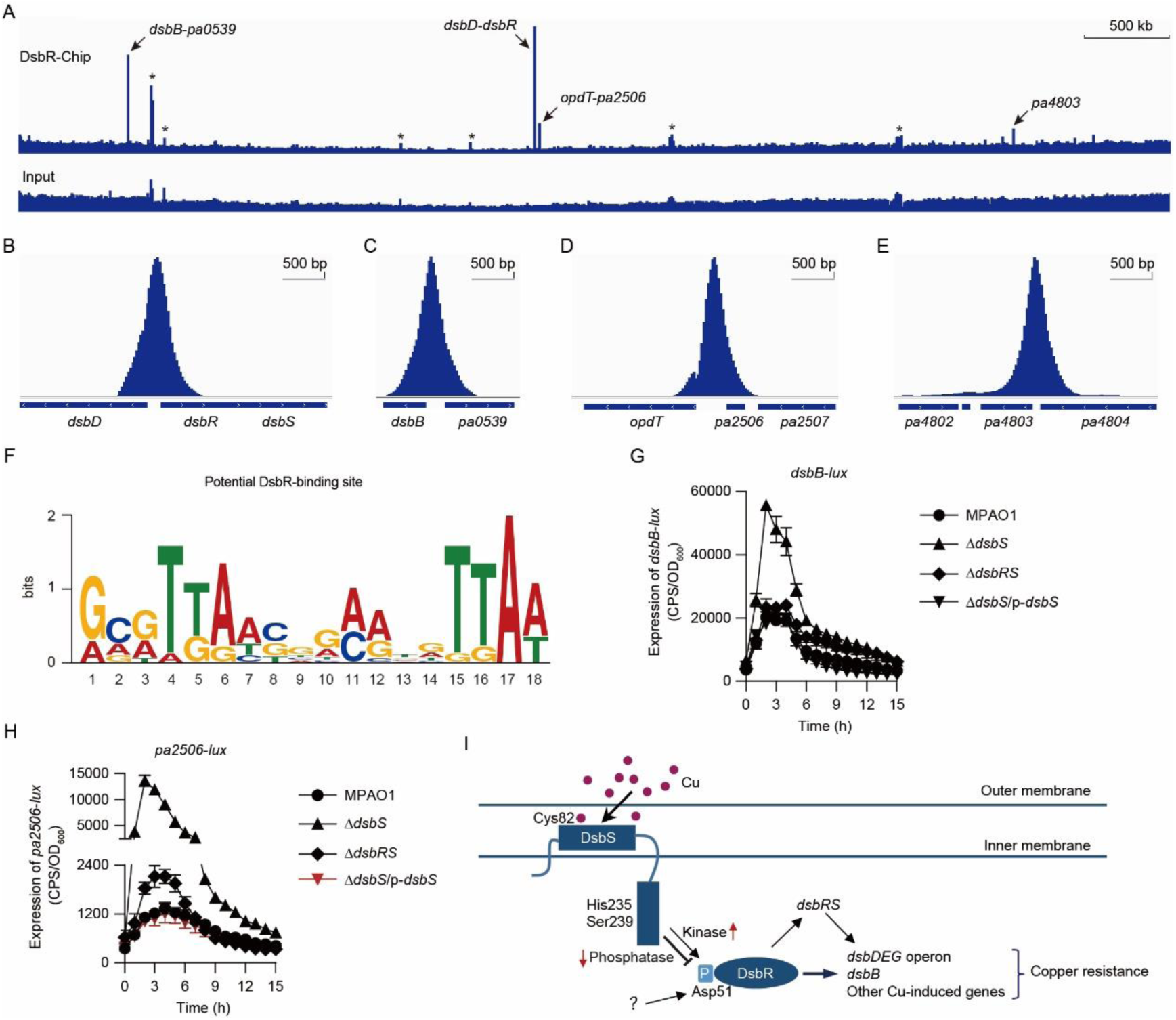
Genome-wide identification of DsbR targets. (A) Representative images of the DsbR CHIP data illustrated by Integrated Genome Viewer software. The y-axis indicates the ChIP-seq signal. The asterisks indicate the false positive peaks, which also identified in the input ChIP-seq experiment that serves as a control. Arrows show the top four ChIP-seq peaks (mean enrichment > 20-fold). (B to E) Schematic presentation of the top four ChIP-seq peaks including those corresponding *dsbD*-*dsbR* intergenic region (B), *dsbB*-*pa0539* intergenic region (C), *opdT*-*pa2506* intergenic region (D), and the promoter region of *pa4803* (E). Values along the y-axis represent the ChIP-seq signal, in arbitrary units. (F) The top-ranking motif identified by MEME program. Motif search was performed on ten 151-bp sequences centered at ten ChIP-seq peaks (mean enrichment > 4-fold) with default parameters except that the site distribution was set to any number of repetitions (anr). The motif was produced (with a E-value of 1.1e-02) in 8 out of 10 input sequences, and the height of each letter represents the relative frequency of each base at different positions in the consensus. (G and H) Expression of *dsbB*-*lux* (G) and *PA2506*-*lux* (H) in *P. aeruginosa* strains cultured in LB. MPAO1, Δ*dsbS*, and Δ*dsbRS* harbor the control plasmid pAK1900; p-*dsbS* denotes pAK1900-*dsbS*. Data from n = 3 biological replicates reported as mean ± SD. (I) A proposal model for mechanism of action of in *P. aeruginosa* DsbRS. In brief, DsbS is a bifunctional histidine kinase, acting as a phosphatase toward phosphorylated DsbR in the absence of copper stimulus. Upon copper binding to its periplasmic domain, the phosphatase activity of DsbS is inhibited, resulting in an activation of DsbR, which in turn triggers the transcription of genes involved in copper stress-response. The lines show the interaction between the players: arrow, activation; hammerheads, repression; solid line, a direct connection. The question mark (“?”) denotes a yet-unidentified effect.

To identify *cis*-regulatory sequences mediating DsbR binding, we performed a *de novo* motif discovery on 10 DsbR-binding sites (≥ average 4-fold enrichment of three biological replicates) detected by CHIP-Seq using the MEME program. We found that the most significant motif containing a consensus TTAA-N_7_-TTAA sequence (where N is any nucleotide) was enriched in the 151 bp region centered at the summits of peaks for 8 DsbR targets (Fig. 7F). Using EMSAs, we found that SUMO-DsbR could bind to either a DNA fragment covering the intergenic regions between *dsbB* and *pa0539* or a DNA fragment covering the intergenic regions between *opdT* and *pa2506*, but not to an unrelated DNA fragment (*dsb*^SD^) that encodes the periplasmic domain of DsbS (Fig. S5B). These results are in good agreement with those obtained from the CHIP-Seq experiments. Additionally, using promoter-*lux* fusion assays, we showed that the promoter activities of *dsbB* and *pa2506* were positively regulated by DsbR (Fig. 7G and H). Thus, in addition to verifying that *dsbDEG* is a major target of DsbR, these results provide a broader view of the cellular processes influenced by DsbRS.

### Cognates of *dsbRS*-*dsbDEG* pair are widely distributed in bacteria

Using a BLASTP search at NCBI, we found that DsbRS homologs (BLASTP, E-value ≤ 1e-5) are widely present in bacteria. Interestingly, our search revealed that *dsbRS* and *dsbD*, *dsbE*, or *dsbG* homolog gene(s) are physically linked on either the chromosomes or the plasmid of a number of bacterial species belong to different phylum of bacteria, including *actinobacteria*, *firmicutes*, and *proteobacteria* (Fig. S6). In most cases, the Dsb genes in these bacterial species appear to form an operon, where *dsbD* is the first gene and is divergently transcribed from the *dsbRS* (Fig. S6).

Because the distribution of bacterial genes is not random and the functionally linked genes are often found in clusters (52), we reasoned that DsbRS might have a role in regulating the Dsb genes in those bacterial species. To test this hypothesis, we performed *de novo* motif discovery on the promoters (140 bp DNA sequence upstream of ATG) of the 14 DsbD genes (Fig. S6) using the MEME program (53). We found that the most statistically significant motif (E-value: 3.0e-39) produced by MEME corresponds to the DNA sequences recognized by DsbR in the intergenic regions between *dsbD* and *DsbR* (Fig. S6, Fig. 1 F and G). These results suggest that regulation of Dsb genes by DsbR might be evolutionarily conserved in bacteria.

Noticeably, Cys82 and the autophosphorylation site of *P. aeruginosa* DsbS (i.e., His235) are completely conserved in all DsbS homologs examined (Fig. S7). Ser239, a residue required for the phosphatase activity of *P. aeruginosa* DsbS, is also conserved, albeit to a lesser extent (Fig. S7). Indeed, DsbS homologs shared amino acid sequence similarity not only in the C-terminal regions but also in the N-terminal regions, where the periplasmic domain of DsbS is located (Fig. S7). In contrast, the amino acid sequence of the periplasmic domain of DsbS showed no significant similarity to those of characterized copper-sensing HKs including CopS, CinS, PcoS, and CusS (Fig. S8), indicating a hitherto unknown Cu-binding mechanism for DsbS. In sum, these data suggest that regulation of Dsb genes by DsbRS may represent an evolutionarily conserved mechanism by which diverse bacteria sense and respond to copper.

## Discussion

The most rapid and efficient means of adapting gene transcription to extracellular stresses often involves TCSs (10–13). Here, we report a novel copper-sensing TCS (designated DsbRS) that confers copper resistance in *P. aeruginosa* (Fig. 7I). We defined the DsbRS regulon consisting of at least 23 genes including those involved protein disulfide bond formation such as *dsbDEG* operon and *dsbB* (Fig. 1, Fig. 7, Data set 1, sheet 1). To our knowledge, this is the first identification of a TCS dedicated to the regulation of Dsb genes (Fig. 7I).

Many genomes of Gram-negative bacteria encode copper-sensing TCS that is responsible for periplasmic copper detoxification (8, 54). The regulons of the canonical copper-sensing TCSs are genes encoding for Cus system (CusRS and CusCFBA), Pco system (PcoABCDRSE), and Cop system (CopABCDRS) that control periplasmic copper homeostasis (11, 54). Interestingly, the DsbRS regulon (Data set 1, sheet 1) differed significantly from those of other characterized copper-sensing TCSs (e.g., CusRS, CopRS, PcoRS, and CinRS), indicative of a distinct mechanism of action of DsbRS. Additionally, complementation of *P. aeruginosa dsbRS* deletion mutant with *dsbDEG* operon, a target of DbsRS (Fig. 1 F and G, Fig. 7A), was able to restore the copper resistance phenotype to a wild-type level (Fig. 6A). Therefore, activation of Dsb genes by DsbRS may represent a novel regulatory axis involved in bacterial copper resistance.

Dsb-family proteins play diverse roles in the conversion between the oxidized and reduced states of cysteine residues of substrate proteins (26-29, 40, 55). Now two distinct pathways for protein disulfide bond formation have been well characterized in *E. coli*, the DsbA-DsbB oxidative pathway, which catalyzes disulfide bond formation, and the DsbC-DsbD reductive pathway, which shuffles incorrectly formed disulfides (26, 29, 55). However, many bacterial genomes encode a repertoire of thiol-disulfide oxidoreductases that is significantly expanded the Dsb system (56, 57). For instance, *P. aeruginosa* has multiple Dsb proteins, including two DsbA proteins [DsbA1 (PA5489) and DsbA2 (PA0982)] and two DsbB proteins [DsbB1 (PA0538) and DsbB2 (PA5256)] involved in the oxidative pathway (58). The expression of either DsbA or DsbB genes in *P. aeruginosa* may be differently modulated according to the environmental conditions (58). Supporting this notion, we showed that *dsbB1* (*pa0538*), but not *dsbB2* (*pa5256*), is a target of DsbRS under our testing conditions (Fig. 7, Data set 1, sheet 1).

We found that the *dsbDEG* operon, encoding proteins potentially involved in the DsbC-DsbD reductive pathway that shuffles incorrectly formed disulfides, is a major target of DsbRS (Fig. 1, Fig. 2, and Fig. 7 A and B). Among them, DsbE and DsbG are disulfide isomerases, like DsbC, and play important roles in the rearrangement of disulfide bonds but with presumably different substrate specificity (59, 60), whereas DsbD is a cytoplasmic membrane protein that maintains these disulfide isomerases in their reduced and thereby active form (26-30, 55, 57, 61). In *P. aeruginosa*, the expression of *dsbDEG* operon is strongly induced by copper (36) and its deletion render the cells to be more susceptible to copper stress conditions (Fig. 6). Thus, it appears that the Dsb system, especially the reductive pathway that ensures isomerization of non-native disulfide bonds, plays an important role in the copper resistance in *P. aeruginosa*.

We showed that in the absence of copper stimulus, DsbS acts as phosphatase to inhibit the phosphorylation and activation of DsbR (Fig. 2, Fig. 3 D and E). For Cu-induced activation of DsbR, not only the reduction of phosphatase activity but also the autokinase of DsbS was crucial (Fig. 3G). Therefore, it is reasonable to assume that when DsbS detects periplasmic copper overload, its phosphatase activity is inhibited, allowing the accumulation of DsbR∼P, a consequence of either Cu-dependent blockage of the phosphatase activity or Cu-dependent stimulation of the autokinase activity of DsbS. However, for unknown reasons we failed to purify the full-length DsbS, and the detailed mechanisms by which the Cu alters the autokinase and phosphatase activities of DsbS await further elucidation. Nonetheless, DsbR can be phosphorylated in the absence of *dsbS* in *P. aeruginosa* (Fig. 2). DsbR can also be phosphorylated by endogenous phosphor-donor acetyl phosphate *in vitro* (Fig. S2 A and B). These observations suggest that inhibition of the phosphatase activity of DsbS might also cause phosphorylation of DsbR by either endogenous phosphor-donors or a non-cognate HK, or both, which in turn contributes to the activation of DsbR (Fig. 7I).

In summary, in this study we identified a novel copper-sensing TCS involved in the copper resistance of *P. aeruginosa*. Further study of the regulation mechanism of DsbRS and its homologs should help us better understand how bacteria respond and adapt to copper stress. A deep knowledge of bacterial copper resistance will ultimately help to develop novel Cu-based prevention and intervention strategies against bacterial infections.

## Materials and Methods

### Bacterial strains and culture conditions

Table S1 lists the bacterial strains and plasmids used in this study. Unless noted otherwise, *Pseudomonas aeruginosa* and *Escherichia coli* were grown in Luria-Bertani (LB) broth. Cultures were incubated at 37°C with shaking (250 rpm). For plasmid maintenance, antibiotics were used at the following concentrations: for *P. aeruginosa*, gentamicin at 30 μg/ml in LB or 150 μg/ml in Pseudomonas Isolation Agar (PIA; BD); tetracycline at 30 μg/ml in LB and or 150 μg/ml in PIA; carbenicillin at 100 μg/ml in LB and PIA; for *E. coli*, carbenicillin at 100 μg/ml, kanamycin at 50 μg/ml, tetracycline at 10 μg/ml, and gentamicin at 10 μg/ml.

### Construction of Δ*dsbS*, Δ*dsbRS*, and Δ*dsbDEG* mutants

For gene replacement, a SacB-based strategy was employed as previously described (46, 62, 63). To construct the *dsbS* null mutant (Δ*dsbS*), polymerase chain reactions (PCRs) were performed to amplify sequences upstream (∼1 kb) and downstream (∼0.9 kb) of the intended deletion. The upstream fragment was generated by PCR using MPAO1 genome as the template and primer pair D-dsbS-up-F/R (*Sac*I/*Sma*I) (Table S2), while the downstream fragment was generated by PCR using primer pair D-dsbS-down-F/R (*Sma*I/*Hin*dIII). The two PCR products were digested and then cloned into *SacI*/*Hin*dIII-digested gene replacement vector pEX18Ap, yielding pEX18Ap::*dsbS*UD. Subsequently, a *ca*. 1.8 kb gentamicin resistance cassette cut from pPS858 with *Sma*I was cloned into pEX18Ap::*dsbS*UD, giving pEX18Ap::*dsbS*UGD. The resultant plasmid was electroporated into wild-type MPAO1 with selection for gentamicin resistance. We chose colonies which show both gentamicin resistance and loss of sucrose (5%) susceptibility, which typically indicates a double-cross-over event and thus of gene replacement occurring. The Δ*dsbS* mutant was further confirmed by PCR.

A similar strategy was used to construct the Δ*dsbRS* mutant. Briefly, a ∼1.1 kb upstream fragment and a ∼0.9 kb upstream fragment of the intended deletion were amplified by PCR using primer pairs D-dsbRS-up-F/R (*Eco*RI/*Bam*HI) and D-dsbRS-down-F/R (*Bam*HI/*Hin*dIII), respectively. The two PCR products were digested and then cloned into *Eco*RI/*Hin*dIII-digested gene replacement vector pEX18Ap, yielding pEX18Ap::*dsbRS*UD. Subsequently, a *ca.* 1.8 kb gentamicin resistance cassette cut from pPS858 with *Bam*HI was cloned into pEX18Ap::*dsbRS*UD, giving pEX18Ap::*dsbRS*UGD. For the construction of Δ*dsbDEG* mutant, a ∼1 kb upstream fragment and a ∼0.9 kb upstream fragment were amplified by PCR using primers D-dsbDEG-up-F/R (*Sac*I/*Kpn*I) and D-dsbDEG-down-F/R (*Kpn*I/*Hin*dIII), respectively. The two PCR products were digested and then cloned into *Sac*I/*Hin*dIII-digested gene replacement vector pEX18Ap, yielding pEX18Ap::*dsbDEG*UD. A *ca.* 1.8 kb gentamicin resistance cassette cut from pPS858 with *Kpn*I was cloned into pEX18Ap::*dsbDEG*UD, giving pEX18Ap::*dsbDEG*UGD.

### Construction of vectors

To construct the plasmid for constitutive expression of *dsbR*, a ∼0.8 kb PCR product covering 112 bp of the *dsbR* upstream region, the *dsbR* gene, and 3 bp of the downstream of *dsbR* was amplified with primer pair *dsbR*-comp-F/R (*Hin*dIII/*Kpn*I). The product was digested with *Hin*dIII and *Kpn*I and ligated into pAK1900 (64) in the same orientation as p*lac* to generate p-*dsbR*. For generating p-*dsbS*, a ∼1.4 kb PCR product covering 13 bp of the *dsbS* upstream region, the *dsbS* gene, and 90 bp of the downstream of *dsbS* was amplified using primer pair *dsbS*-comp-F/R (*Hin*dIII/*Bam*HI). For generating p-*dsbRS*, a ∼2.2 kb PCR product covering 63 bp of the *dsbR* upstream region, the *dsbRS* operon, and 34 bp downstream of the *dsbS* was amplified using primer pair *dsbRS*-comp-F/R (*Hin*dIII/*Xba*I). For generating p-*dsbDEG*, a ∼3.4 kb PCR product covering 50 bp of the *dsbD* upstream region, the *dsbDEG* operon, 50 bp of the downstream of *dsbG* was amplified using primer pair dsbDEG-comp-F/R (*Bam*HI/*Sac*I). Primers are listed in Table S2 and genomic DNA from *P. aeruginosa* MPAO1 were used as a template for the PCR. All the PCR products were digested with corresponding enzymes and cloned into pAK1900, and the constructs were sequenced in order to ensure that no unwanted mutations resulted.

### Construction for the expression of fusion proteins in *P. aeruginosa*

For generating p-*dsbR*-*flag*, a ∼0.8 kb PCR product covering 14 bp upstream and the *dsbR* gene (not including the stop codon) was generated from MPAO1 genomic DNA with primer pair Flag-dsbR-F/R (*Hin*dIII/*Kpn*I). The *Hin*dIII and *Kpn*I digested PCR product was cloned into pRK415 (65) to generate p-*dsbR*-*flag*. For generating p-*dsbS*-*flag*, a ∼1.3 kb PCR product covering 9 bp upstream and the *dsbS* gene (not including the stop codon) was generated from MPAO1 genomic DNA with primers Flag-dsbS-F/R (*Hin*dIII/*Kpn*I). The *Hin*dIII and *Kpn*I digested PCR product was cloned into pAK1900 to generate p-*dsbS*-*flag*.

p-*dsbR*(D51A)-*flag*, p-*dsbS*(C82A)-*flag*, p-*dsbS*(H235A)-*flag*, and p-*dsbS*(S239A)- *flag* were obtained using the QuikChange II site-directed mutagenesis kit (Stratagene, catalog^#^: 200518). For generating p-*dsbR*(D51A)-*flag*, the primer pair dsbR(D51A)-F/R was used. For generating p-*dsbS*(C82A)-*flag*, primer pair dsbS(C82A)-F/R was used. For generating p-*dsbS*(H235A)-*flag*, primer pair dsbS(H235A)-F/R was used. For generating p-*dsbS*(S239A)-*flag*, primer pair dsbS(S239A)-F/R was used.

For generating p-*dsbS*^Del-SD^-*flag*, we synthetized the DNA fragment covering 9 bp upstream and the truncated *dsbS* (with deletion of sequence encoding the periplasmic domain of DsbS, i.e., residues 28-146) and cloned it into pUC57 vector, yielding pUC57:: *dsbS*^Del-SD^. Subsequently, a ∼1 kb corresponding PCR product was generated from pUC57:: *dsbS*^Del-SD^ with primer pair Flag-dsbS-F/R (*Hin*dIII/*Kpn*I). The *Hin*dIII and *Kpn*I digested PCR product was cloned into pAK1900 to generate p-*dsbS*^Del-SD^-*flag*. All constructs were sequenced to ensure that no unwanted mutations resulted.

### Monitoring gene expression by *lux*-based reporters

The plasmids mini-CTX-*lux* (66) was used to construct promoter-*luxCDABE* reporter fusions *dsbD*-*lux*, *dsbR*-*lux, dsbB*-*lux*, and *pa2506*-*lux*. For *dsbD*-*lux*, the *dsbD* promoter region (−531 to +272 of the start codon) was amplified by PCR using the primer pair mini-dsbD-pro-F/R (*Hin*dIII/*Eco*RI) and cloned into the *Hin*dIII and *Eco*RI sites of the mini-CTX-*lux* to generate mini-CTX-*dsbD-lux*. For *dsbR*-*lux*, the *dsbR* promoter region (−480 to +122 of the start codon) was amplified by PCR using the primer pair mini-dsbR-pro-F/R (*Eco*RI/*Bam*HI) and cloned into the *Eco*RI and *Bam*HI sites of the mini-CTX-*lux* to generate mini-CTX-*dsbR-lux*. For *dsbB*-*lux*, the *dsbB* promoter region (−235 to +140 of the start codon) was amplified by PCR using the primer pair mini-dsbB-pro-F/R (*Hin*dIII/*Bam*HI) and cloned into the *Hin*dIII and *Bam*HI sites of the mini-CTX-*lux* to generate mini-CTX-*dsbB*-*lux*. For *pa2506*-*lux*, the *pa2506* promoter region (−396 to +106 of the start codon) was amplified by PCR using the primers mini-*PA2506*-pro-F/R (*Hin*dIII/*Bam*HI) and cloned into the *Hin*dIII and *Bam*HI sites of the mini-CTX-*lux* (Table S1) to generate mini-CTX-*pa2506-lux*. All the promoters are oriented in the same direction as *luxCDABE*, and constructs were sequenced to ensure that no unwanted mutations resulted. The resulting plasmid was conjugated into *P. aeruginosa* strains and the construct was integrated into the *attB* site as described previously (46, 62, 63) though a diparental mating using *E. coli* S17 λ-pir as the donor. In MPAO1 and its derivatives, parts of the mini-CTX-*lux* vector containing the tetracycline resistance cassette were deleted using a flippase (FLP) recombinase encoded on the pFLP2 plasmid (46, 62, 63).

Unless noted otherwise, the expression of promoter fusion genes was measured in a 96-well black-wall clear-bottom plate (Corning incorporated, Costar, Code^#^: 3603) as previously described (46, 62). Briefly, overnight LB cultures were diluted to an OD_600_ of 0.05 in LB medium, and then a 100 μl volume of the sample was added to the wells, and subsequently a 50 μl volume of filter-sterilized mineral oil was added in order to prevent evaporation during the assay. Promoter activities were measured as counts per second (CPS) of light production with a Synergy 2 Multi-Mode Microplate.

### Analysis of *dsbDEG* transcripts

Overnight culture of the Δ*dsbS* strain was washed and diluted 100-fold in LB medium (OD_600_ ≈ 0.05). 20 ml liquid cultures were grown in a 100 ml flask at 37 °C, shaking with 250 rpm for 3 h. Total RNA was immediately stabilized with RNA protect Bacteria Reagent (Qiagen, CAS^#^:76506) and then extracted by using a Qiagen RNeasy kit (Qiagen, catalog^#^: 74104). The total DNase-treated RNA (5 μg) was reversely transcribed to synthesize cDNA using the Hifair® II 1st Strand cDNA Synthesis Kit (YEASEN, Lot:11123ES10) with random primers according to the manufacture’s recommendation. Amplification of DNA fragment covering +1477 (relative to start codon) of *dsbD*, *dsbE*, and +190 of *dsbG* was done using primer pair RT-*dsbDEG*-F/R (Table S1). The amplification products were analyzed by agarose gel electrophoresis and stained with ethidium bromide, and the PCR products were subjected to sequencing to verify the RT-PCR results.

### Overexpression of recombinant proteins in *E. coli* and their purifications

The following six recombinant proteins were expressed in *E. coli*: 1) SUMO-DsbR, the N-terminal 6His-SUMO-tagged DsbR; 2) DsbS^CTD^, the N-terminal 6His-tagged cytosolic segment of DsbS^CTD^ (residues 225-440); 3) DsbS^CTD^H235A, a DsbS^CTD^ mutant in which the histidine 235 (H235) was replaced by alanine (A); 4) DsbS^CTD^S239, a DsbS^CTD^ mutant in which the serine 239 (S239) was replaced by alanine (A); 5) SUMO-DsbS^SD^, the N-terminal 6His-SUMO-tagged sensor domain of DsbS (residues 28-146); 6) SUMO-DsbS^SD^C82A, a SUMO-DsbS^SD^ mutant in which the cysteine (C82) was replaced by alanine (A).

For the construction of expression plasmid of the SUMO-DsbR proteins, primer pair pET-dsbR-F/R (*Sac*I/*Hin*dIII) was used. The DNA fragment amplified from *P*. *aeruginosa* MPAO1 genomic DNA encoding the full-length DsbR was cloned into pET28a(+)-sumo, yielding pET28a(+)-sumo::*dsbR* plasmid. For the construction of the expression plasmid of DsbS^CTD^ proteins, primer pair pET-CTD-F/R (*Nde*I/*Bam*HI) was used and the corresponding DNA fragment encoding the kinase domain of DsbS (residues 225-440) was cloned into the vector pET28a (Table S1), yielding pET28a::*dsbS*^CTD^ plasmid. For the construction of the expression plasmid of SUMO-DsbS^SD^ proteins, the DNA sequence of the sensor domain of DsbS (residues 28-146) was amplified from *P*. *aeruginosa* MPAO1 genomic DNA with the primer pair pET-SD- F/R (*Sac*I/*Hin*dIII) by PCR and subsequently cloned into pET28a(+)-sumo, yielding pET28a(+)-sumo::*dsbS*^SD^ plasmid for the expression of SUMO-DsbS^SD^ proteins. For generating pET28a::*dsbS*^CTD^H235A, the primer pair dsbS(H235A)-F/R and a QuikChange II site-directed mutagenesis kit (Stratagene, catalog^#^:200518) were used. For generating pET28a::*dsbS*^CTD^S239A, the primer pair dsbS(S239A)-F/R and a QuikChange II site-directed mutagenesis kit were used. For generating pET28a(+)- sumo::*dsbS*^SD^C82A, the primer pair dsbS(C82A)-F/R and a QuikChange II site-directed mutagenesis kit were used. All constructs were sequenced to ensure that no unwanted mutations resulted.

The protein was expressed in *E. coli* strain BL21 star (DE3) and purifications were performed as previously described (67, 68). Briefly, bacteria were grown at 37°C overnight in 10 ml of LB medium with shaking (250 rpm). The cultures were transferred into 1 L of LB medium (containing 50 µg/ml kanamycin) incubated at 37°C with shaking (200 rpm) until the OD_600_ reached 0.6, and then IPTG (isopropyl-1-thio-β-d-galactopyranoside) was added to a final concentration of 1.0 mM. After 20 h incubation at 16°C with shaking (200 rpm), the cells were harvested by centrifugation and stored at −80°C. The cells were lysed at 4°C by sonication in buffer A (50 mM Tris, pH 8.0; 250 mM NaCl; 1 mM DTT and 20 mM imidazole). The whole cell fraction was subjected to centrifugation at 4°C at 12, 000 rpm for 25 min to remove insoluble material and the membrane fraction. The supernatant from this pellet was loaded onto a HisTrap HP column (GE Healthcare, Lot: 10241151), equilibrated with buffer A and eluted with a 0-100% gradient of buffer B (20 mM Tris-HCl, pH 8.0; 250 mM NaCl; 1 mM DTT; 500 mM imidazole). The fractions containing proteins were loaded onto the HiTrap Desalting 5 x 5 ml (Sephadex G-25 S) (GE Healthcare, Code^#^: 17-1408-01) with a running condition of 50 mM Tris (pH 8.0), 250 mM NaCl and 1 mM DTT to remove the imidazole. The purified protein was > 90% pure as estimated by a 12% (wt/vol) SDS/PAGE gel.

### Cleavage and removal of SUMO tag in SUMO-DsbS^SD^ and SUMO-DsbS^SD^C82A

Add the following to a 15 ml centrifuge tube: the fusion Protein 1 mg, 10 X SUMO protease Buffer (500 mM Tris-HCl, pH 8.0; 2 M NaCl; 50% glycerol; 10 mM DTT) 1 ml, ddH_2_O 8.5 ml, SUMO protease (Introvogen, catalog^#^: 12588018) (10 units) 500 μl then incubate at 4°C overnight. Analyze 30 μl of sample by SDS-PAGE using a 15% SDS/PAGE gel and determine the percent protein cleavage by analyzing the amount of cleaved products formed and amount of uncleaved protein remaining after digestion.

The SUMO protease and SUMO tag both contain a polyhistidine tag at the N-terminus. So after cleavage of the fusion protein, the removal of SUMO protease and SUMO tag from the cleavage reaction was performed with affinity chromatography on a HisTrap HP column (GE Healthcare, Code^#^:17-5248-01). The SUMO protease and SUMO tag will stay in the HisTrap HP column and the cleaved native protein will be in the flow-through fractions. The purified Dsb^SD^ or Dsb^SD^C82A was > 95% pure as estimated by a 15% SDS/PAGE gel.

### Electrophoretic mobility shift assay (EMSA)

The electrophoretic mobility shift experiments were performed as previously described (46, 67). Briefly, 20 µl of the DNA probe mixture (30 to 50 ng) and various amounts of purified proteins in binding buffer (10 mM Tris-Cl, pH 8.0; 1 mM DTT; 10% glycerol; 5 mM MgCl2; 10 mM KCl) were incubated for 30 min at 37°C. When indicated, either 30 mM acetyl phosphate or 0.5 μl Calf Intestinal Alkaline Phosphatase (CIP) (NEB, catalog^#^: M0290V) was added to the solution. Native polyacrylamide gel (6%) was run in 0.5 × TBE buffer at 90 V at 4°C. The gel was stained with GelRed nucleic acid staining solution (Biotium) for 10 min, and then the DNA bands were visualized by gel exposure to 260-nm UV light.

DNA probes were PCR-amplified from *P. aeruginosa* MPAO1 genomic DNA using the primers listed in Table S2. For *dsbD* promoter, a 578 bp DNA fragment covering the promoter region of *dsbD* (from −550 to +28 of the start codon) was amplified using primer pair EMSA-dsbD-pro-F/R. For *dsbB* promoter, a 475 bp DNA fragment covering the promoter region of *dsbB* (from −381 to +94 of the start codon) was amplified using primer pair EMSA-dsbB-pro-F/R. For *PA2506* promoter, a 478 bp DNA fragment covering the promoter region of *PA2506* (from −376 to +102 of the start codon) was amplified using primer pair EMSA-PA2506-pro-F/R. All PCR products were purified by using a QIAquick gel purification kit (QIAGEN, catalog^#^: 28104).

### Dye primer-based DNase I footprinting assay

The published DNase I footprint were performed as previously described (46, 67). Briefly, PCR was used to generate DNA fragments using the primers dsbR-pro-F and dsbR-pro-R-FAM. PCR products were purified by with QIAquick gel purification kit. Subsequently, 50 μl reaction mixture containing 300 ng 6-carboxyfluorescein (6-FAM)-labeled promoter DNA, 3.4 μM (or indicated) of SUMO-DsbR, followed by the addition of binding buffer (10 mM Tris-Cl, pH 8.0; 1 mM DTT; 10% glycerol; 1 mM EDTA; 50 mM NaCl) up to 50 μl and incubated at room temperature for 30 min. Subsequently, 0.01 Unit of DNase I (Promega Biotech Co., Ltd, catalog^#^: 137017) was added and the reaction mixture incubated for additional 5 minutes. The DNase I-digestion was terminated by adding 90 μl of quenching solution (200 mM NaCl, 30 mM EDTA, 1% SDS), and then the mixture was extracted with 200 μl of phenol-chloroform-isoamyl alcohol (25:24:1). The digested DNA fragments were isolated by ethanol precipitation, dried under vacuum and resuspended in RNase-free water. Then, 5 μl of digested DNA was mixed with 4.9 μl of HiDi formamide and 0.1 μl of GeneScan-500 LIZ size standards (Applied Biosystems). A 3730xl DNA analyzer was used to detect the sample, and the result was analyzed with GeneMapper software (Applied Biosystems). The dye primer based Thermo SequenaseTM Dye Primer Manual Cycle Sequencing Kit (Thermo, Lot:4313199) was used in order to more precisely determine the sequences of the DsbR-protection region, and the corresponding label-free promoter DNA fragment was used as template for DNA sequencing. Electropherograms were analyzed with GeneMarker v1.8 (Applied Biosystems).

### Immobilized metal ion affinity chromatography (IMAC)

IMAC was performed as previously described (21). Briefly, a 100-μL aliquot of Hisbind resin (Novagen, catalog^#^: 69670) was loaded with 500 μl of 0.5 mM CuSO_4_, NiSO_4_, ZnSO_4_, or CoCl_2_ in water and then equilibrated in buffer R (50 mM Tris-HCl, pH 8.0; 250 mM NaCl). About 10 μg of purified DsbS^SD^ were applied to the columns. After centrifugation, the supernatants (flowthrough fractions) were removed and the columns were washed three times with 1 ml of buffer R. The proteins were then eluted with 100 μl of 0.5 M imidazole in buffer R. Fifteen microliters of flowthrough fractions and the imidazole eluted were analyzed by 15% SDS-PAGE and Coomassie Blue staining.

### Isothermal titration calorimetry

Microcalorimetric titrations were carried out at 25°C using a MicroCal ITC200 (Malvern). After removal of SUMO tag in SUMO-DsbS^SD^, the fractions containing DsbS^SD^ were concentrated and loaded onto a Superdex-75 gel filtration column with a running condition of 50 mM Tris-HCl, pH 8.0, 250 mM NaCl to remove the glycerol. The concentrations of DsbS^SD^ were determined by the A280 as measured by a Nanodrop2000 (Thermo Scientific). CuSO_4_ was prepared in the same buffer containing 50 mM Tris-HCl, pH 8.0, 250 mM NaCl. 20 mM CuSO4 was injected 20 times (1 μl for injection 1 and 2 μl 101 for injections 2–20), with 120 s intervals between injections. Raw titration data were concentration-normalized and corrected for dilution effects prior to analysis using the “One-binding site model” of the MicroCal version of ORIGIN. The parameters Δ*H* (reaction enthalpy), *K*_A_ (binding constant, *K*_A_ = 1/*K*_D_) and n (reaction stoichiometry) were determined from the curve fit. The changes in free energy (Δ*G*) and entropy (Δ*S*) were calculated from the values of *K*_A_ and Δ*H* with the equation: Δ*G* = −*RT* ln *K*_A_ = Δ*H* − *T*Δ*S*, where R is the universal molar gas constant and T is the absolute temperature (69).

### Pro-Q Diamond phosphoprotein gel stain

The Pro-Q Diamond phosphoprotein gel stain was performed as previously described with some modifications (46). ∼5 μM SUMO-DsbR was incubated with phosphorylation buffer (10 mM Tris, pH 8.0; 1 mM DTT; 10 mM MgCl_2_; 20 mM KCl) in a final reaction volume of 50 μl at 37°C for 30 min. When indicated, 30 mM acetyl phosphate or 1 µl Calf Intestinal Alkaline Phosphatase (CIP) (NEB, catalog^#^: M0290V) was added to the solution. Aliquots were resolved on a 12% SDS polyacrylamide gel. The phosphorylation of SUMO-DsbR within the gel was detected by using a Pro-Q™ Diamond Phosphoprotein Gel Staining Kit (Invitrogen, catalog^#^: MPP33300) following the manufacturer’s instructions. Fluorescent output was recorded using an Tanon-5200 multi. Protein phosphorylation level was quantified by determining the ratio of the intensity of phosphoprotein in Pro-Q Diamond image to its intensity of total protein in Coomassie blue image using ImageQuant software (Molecular Dynamics, Sunnyvale, CA).

### *In vitro* phosphorylation assay using radiolabelled ATP

Autophosphorylation assays were carried out in autophosphorylation buffer [50 mM Tris-HCl, pH 8.0; 200 mM KCl; 2 mM MgCl_2;_ 0.1 mM EDTA; 10% (vol/vol) glycerol; 10 mM DTT] at room temperature (10). The autophosphorylation was performed on ice with ∼10 µM purified DsbS^CTD^, DsbS^CTD^H235A, or DsbS^CTD^S235A in a final reaction volume of 100 μl. Reactions were started by adding 10 μCi of [γ-^32^P]ATP (PerkinElmer, BLU002A500UC), and 10 μl samples were removed at different times. The reaction was stopped by the addition of 2.5 μl 5 x sodium dodecylsulphate (SDS) loading buffer.

To examine the transphosphorylation of SUMO-DsbR, ∼10 µM DsbS^CTD^ or DsbS^CTD^S235A was incubated with 10 μCi of [γ-^32^P]ATP in the autophosphorylation buffer for 15 min for autophosphorylation at room temperature. SUMO-DsbR at a final concentration of ∼5 μM was then added to the reaction mixture in a final volume of 100 μl. 10 μl aliquots were removed at regular intervals, in which the transphosphorylation reaction was stopped by adding 2.5 μl of 5 × SDS loading buffer.

To examine the dephosphorylation of P∼SUMO-DsbR, ∼5 μM SUMO-DsbR was first transphosphorylated by ∼5 µM phosphorylated DsbS^CTD^S235A for 5 min at room temperature. Then ∼10 µM purified DsbS^CTD^ or DsbS^CTD^S239A was added to 100 μl reaction mixtures in a final volume of 110 μl (DsbS^CTD^/SUMO-DsbR or DsbS^CTD^S239A/SUMO-DsbR ≈ 2:1). Autophosphorylation buffer (10 μl) was also added to 100 μl reaction mixtures as a control. 10 μl aliquots were removed at regular intervals, in which the reaction was stopped by adding 2.5 μl of 5 × SDS loading buffer.

All the samples were analyzed by 12% SDS-PAGE at 4°C, visualized by autoradiography and Coomassie blue staining. Where appropriate, protein phosphorylation level was quantified by determining the ratio of the intensity of autoradiography image to its intensity of total protein in Coomassie blue image using ImageQuant software.

### Western Blot Analysis

Samples resolved on gels were transferred to PVDF (Bio-Rad) membranes through semi-dry transfer assembly (Bio-Rad) for 30 min at room temperature. The membrane was incubated with the primary antibody in 10 ml of 5% (wt/vol) skim milk at 4°C overnight following the blocking step [10 ml of 5% (wt/vol) skim milk] at room temperature for 2 h, and then washed three times at room temperature for 15 min in TBST buffer (10 mM Tris-HCl, pH 7.5; 150 mM NaCl; and 0.1% Tween 20). Then, membranes were incubated with the secondary antibody for 2 h at room temperature and washed three times for 15 min in TBST buffer.

The proteins with Flag tag were detected by Western blot analysis using a mouse anti-Flag monoclonal antibody (Aogma, catalog^#^: AGM12165) followed by a secondary, sheep anti-mouse IgG antibody conjugated to horseradish peroxidase (HRP) (GE Healthcare, Code^#^: NA931). For detection of ClpP, anti-ClpP polyclonal antibody (62) and an anti-rabbit IgG antibody conjugated to horseradish peroxidase (HRP) (GE Healthcare, Code^#^: NA934) were used. For detection of RNA polymerase (RNAP), anti-RNAP (Neoclone, #WP003) antibody and antimouse IgG antibody conjugated to horseradish peroxidase (HRP) (GE Healthcare, Code^#^: NA931). Immunoblots for ClpP or RNAP served as loading control. The images were taken using Tanon-5200 Multi (Tanon, Shanghai, China), according to the manufacturer’s recommendation.

### Sample preparation for *in vivo* detection of DsbR phosphorylation and Phos-tag gel electrophoresis

Overnight LB cultures of *P. aeruginosa* were diluted 100-fold (OD_600_ ≈ 0.05) with fresh LB broth and then the diluted cultures were grown in a 20 ml tube with a flask volume-to-medium volume ratio of 5:1, shaking with 250 rpm. After incubated at 37°C for 4 h (OD_600_ ≈ 2), 100 µl bacteria cells were harvested and immediately resuspended in 60 µl of lysis buffer (50 mM Tris-Cl, pH 7.5; 150 mM NaCl; 1 mM MgCl2; 0.1% Triton X-100; 15 mg/ml DNaseI; 0.5 mM PMSF; 1 mM DTT) with 0.1% (vol/vol) Lysonase (46). Sufficient lysis was achieved by repeated pipetting up and down for 10 s followed by addition of 15 µL of 5 × SDS loading buffer. Samples were immediately loaded onto a Phos-tag gel for electrophoresis (10% acrylamide gels containing 25 µM acrylamide-Phos-tag ligand and 50 µM MnCl_2_) (70). Electrophoresis was performed at 30 mA at 4°C for 2.5 h in Tris-Glycine-SDS running buffer (25 mM Tris, 192 mM glycine, 0.1% SDS, pH 8.4). After electrophoresis, the Phos-tag gel was washed 10 min at room temperature with Transfer Buffer [20%(v/v) methanol, 50 mM Tris, 40 mM glycine] supplied with 1 mM EDTA to remove Mn^2+^ from the gel, then the gel was incubated at room temperature with gentle shaking for another 10 min in Transfer Buffer twice to remove EDTA and followed by Western blot analysis.

### Bacterial growth assays

Bacterial growth in liquid medium was monitored in in a 96-well black-wall clear-bottom plate (Corning incorporated, Costar, Code^#^: 3603) using Synergy 2 Multi-Mode Microplate Reader. Overnight LB cultures of *P. aeruginosa* strains were diluted to an OD_600_ of 0.05 with fresh LB broth. Each well contained 100 μl of culture supplemented without or with CuSO_4_, as indicated. A 50 μl volume of filter-sterilized mineral oil was subsequently added in order to prevent evaporation during the assay. The microtiter plate was incubated at 37°C with gentle shaking for 2 s every 60 min. Growth was monitored by reading the OD_600_ every hour using a Synergy 2 Multi-Mode Microplate Reader.

### Sample preparation for DsbR ChIP-Seq

To perform chromatin immunoprecipitation (ChIP), we firstly constructed chromosomal-borne DsbR-Flag strain Δ*dsbRS*::*dsbR*-*flag*. A ∼1 kb PCR product covering 292 bp of the *dsbR* upstream region and the *dsbR* gene (not including the stop codon) was amplified using primers mini-dsbR-flag-F/R (*Hin*dIII/*Eco*RI). The *Hin*dIII and *Eco*RI-digested PCR product was cloned into the *Hin*dIII and *Eco*RI sites of the mini-CTX-*lux* to generate mini-CTX-DsbR-Flag. The resulting plasmid was conjugated into *P. aeruginosa* Δ*dsbRS* strain and the construct was integrated into the attB site as described previously (46, 62) though a diparental mating using *E. coli* S17 λ-pir as the donor, yielding a Δ*dsbRS*::*dsbR*-*flag* strain (Table S1).

Overnight LB cultures of Δ*dsbRS*::*dsbR*-*flag* were diluted 100-fold (OD_600_ ≈ 0.05) with fresh LB broth and then the diluted cultures were grown in a 250 ml Erlenmeyer flask with a flask volume-to-medium volume ratio of 5:1, shaking with 250 rpm. After incubated at 37°C for 3 h, *P. aeruginosa* cultures were treated with 1% formaldehyde for 10 min at 37°C. Crosslinking was stopped by addition of 125 mM glycine. Bacterial pellets were washed twice with a Tris buffer (20 mM Tris-HCl pH 7.5, 150 mM NaCl), and then resuspended in 500 µl immunoprecipitation buffer [50 mM HEPES-KOH pH 7.5, 150 mM NaCl, 1 mM EDTA, 1% Triton X-100, 0.1% sodium deoxycholate, 0.1% SDS, mini-protease inhibitor cocktail (Roche, catalog^#^: 11 836153001)] and the DNA was sonicated (JY92-IIDN Ultrasonic Homogenizer) to sizes of 100–300 bp (15% total output, 1-second on, 2-second off, for 30 min on ice). Insoluble cellular debris was removed by centrifugation and the supernatant was used in immunoprecipitation (IP) experiments. 300 µl supernatant was added into protein A beads (Smart-Lifesciences, catalog^#^: SA015C), and then incubated with 50 µl agarose-conjugated anti-Fag antibodies (Aogma, catalog^#^: AGM12165) in immunoprecipitation buffer. Washing, crosslink reversal and purification of the ChIP DNA were conducted by following previously published protocol (71). DNA in untreated supernatant was collected as an input sample.

### ChIP-Seq library construction, DNA sequencing, and data analysis

DNA fragments (150–250 bp) were selected for library construction and sequencing libraries prepared using the NEBNext ChIP-Seq Kit (NEB, catalog#: E6200). The libraries were sequenced using the HiSeq XTen (Illumina). Three independent ChIP-Seq experiments were performed and ChIP-seq reads were mapped to the *P. aeruginosa* MPAO1 genomes, using TopHat (Version 2.0.0) with two mismatches allowed (72). Only the uniquely mapped reads were kept for the subsequent analyses. The enriched peaks were identified using MACS software (version 2.1.2) (73) and were visualized and screenshots prepared using Integrative Genome Viewer (IGV) (version 2.4.19) (74). We filtered peaks called by MACS by requiring an adjusted score (i.e., -log10 P-value) of at least 50 in order to ensure that we had a high quality peak annotation. Peaks occurred in all three ChIP-Seq experiments with a fold enrichment ≥ 2 were shown in Data set 1 for each biological replicates (Sheet 2-4). All Chip-seq data (input sample and three independent biological replicates) have been submitted to Gene Expression Omnibus (GEO) under the GEO Series accession number GSE154248, with the following BioSample accession numbers: SAMN15508106 to SAMN15508109. MEME (53) was used for motif discovery on DNA sequences around individual ChIP-seq “peaks” from the ChIP-seq experiments.

## Acknowledgements

This work was supported by grants from the Ministry of Science and Technology (MOST) of China (grant no. 2016YFA0501503 and 2019ZX09721001-004-003) and National Natural Science Foundation (NSFC) (grant nos. 31670136, 31870127, and 81861138047). This study was also funded by the Science and Technology Commission of Shanghai Municipality (grant no. 19JC1416400) and State Key Laboratory of Drug Research (SIMM2003ZZ-03). We appreciate ChIP-seq assays and the corresponding initial data analysis from Sangon Biotech (shanghai) Co., Ltd.

## Author contributions

L.L. and L.Y. conceived and initiated the study. L.Y. did most experiments. Q.C. did Phos-tag assay and performed ITC experiment. L.Y., Q.C., W.C., N.Y., C-G.Y., Q.J., M.W., T.B. and L.L. analysed the data. L.L. supervised the study and wrote the manuscript with input from L.Y., Q.C., and T.B. All authors discussed the results and commented on the manuscript.

## Supplementary Materials

### Figures and Figure legends

**Fig. S1.**
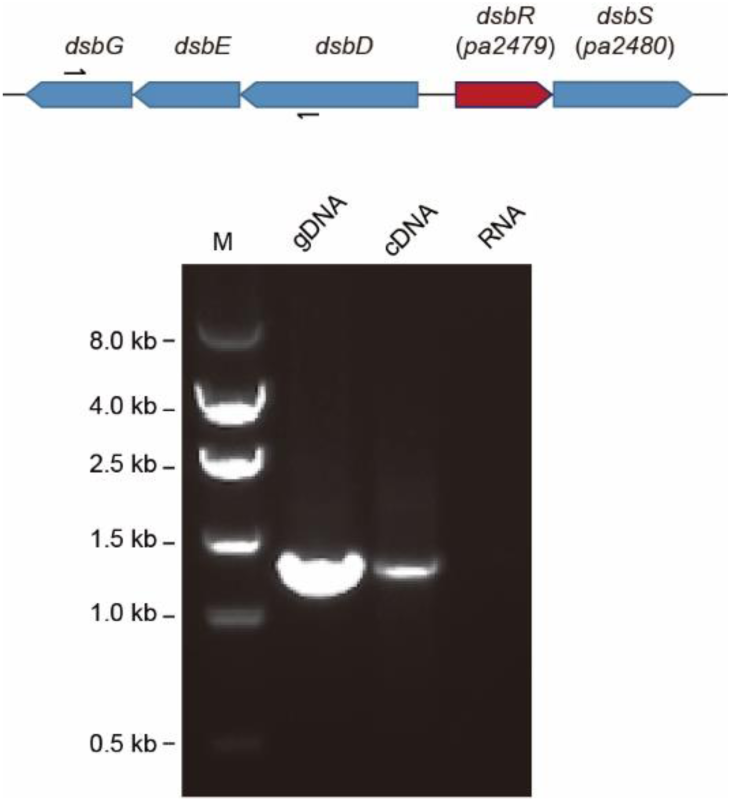
Verification of the operon structure for *dsbG*, *dsbE* and *dsbD* by reverse transcription-PCR (RT-PCR). Upper panel, scheme showing the position and orientation of the primers (black arrows). Lower panel, ethidium bromide-stained agarose gel (1%) of genome-based PCR (gDNA) and whole-cell RNA-based RT-PCR (cDNA) products amplified with primer pair *dsbDEG*-F and *dsbDEG*-R (Table S2); controls without reverse transcriptase added (RNA) were included; M, DNA size standards (in base pairs).

**Fig. S2.**
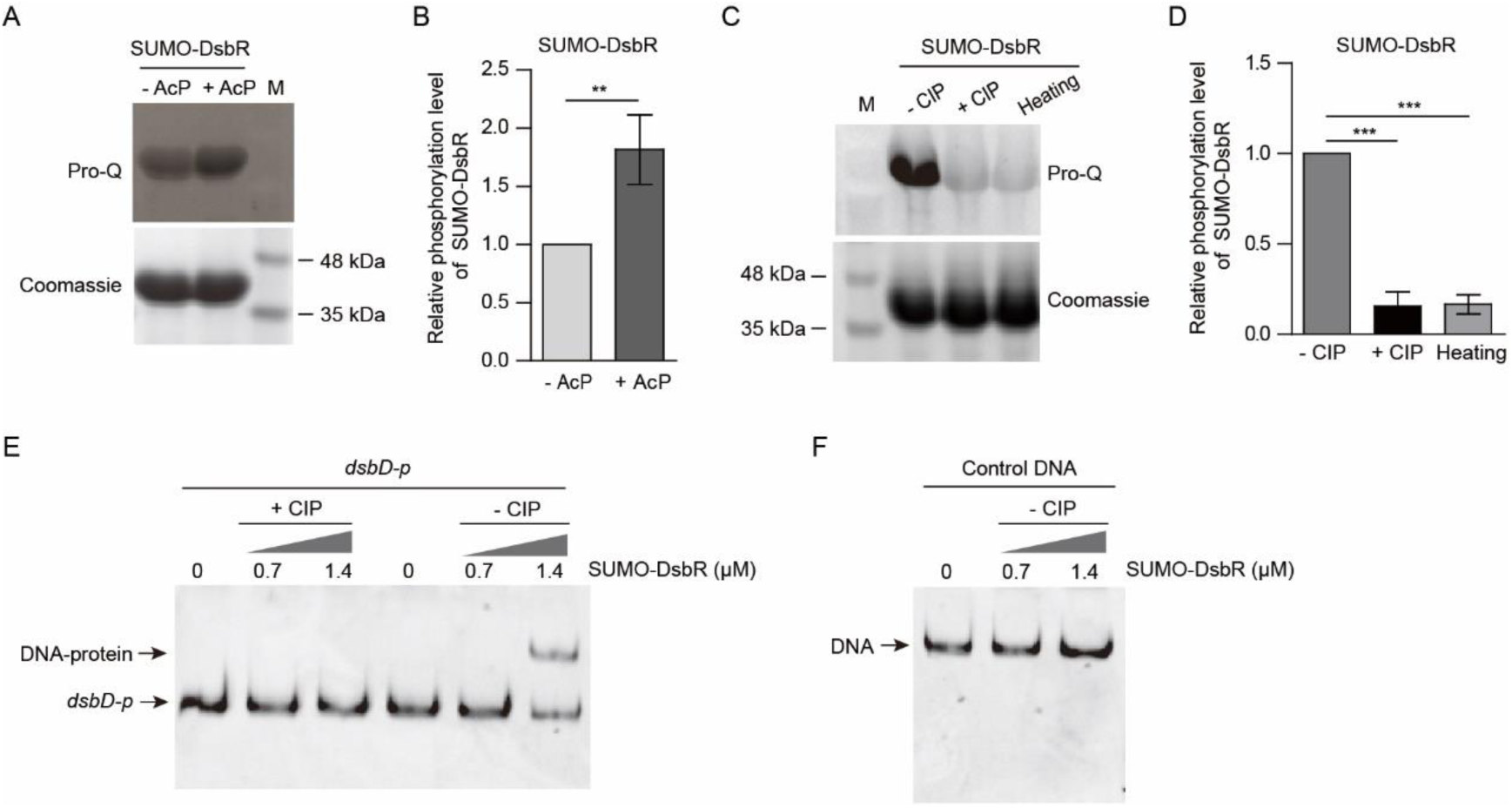
*In vitro* phosphorylation assays and EMSAs. (A and B) Representative images (A) and analysis (B) of the phosphorylation of N-terminal His_6_-SUMO-tagged DsbR (i.e., SUMO-DsbR) in the absence (-) or presence (+) of 30 mM acetyl phosphate (AcP). The intensities of Pro-Q Diamond stained bands of SUMO-DsbR proteins were standardized against the intensities of the same bands restained with Coomassie blue and was further normalized to the vehicle group (-AcP). M, protein marker. Data from n = 3 technical replicates reported as mean ± SD (**p < 0.01, Student’s two-tailed *t-*test). (C and D) Representative images (C) and analysis (D) of the phosphorylation of purified SUMO-DsbR treated without (- CIP) or with calf intestinal alkaline phosphatase (+ CIP), or heating at 95°C for 5 min (Heating). The intensities of Pro-Q Diamond stained bands of SUMO-DsbR proteins were standardized against the intensities of the same bands restained with Coomassie blue and was further normalized to the vehicle group (-CIP). Data from n = 3 technical replicates reported as mean ± SD (***p < 0.001, Student’s two-tailed t-test). (E and F) EMSAs show the DNA-binding ability of purified SUMO-DsbR to the *dsbD*-*dsbR* intergenic region DNA (i.e., *dsbD*-*p*) (C) and a control DNA fragment (D). The purified SUMO-DsbR was pretreated without (- CIP) or with calf intestinal alkaline phosphatase (+ CIP) at 37°C for 30 min. *dsbD*-*p*, a 578-bp DNA covering the intergenic region of *dsbD*-*dsbR*; control DNA, a 673-bp DNA fragment encoding the cytoplasmic domain of DsbS.

**Fig. S3.**
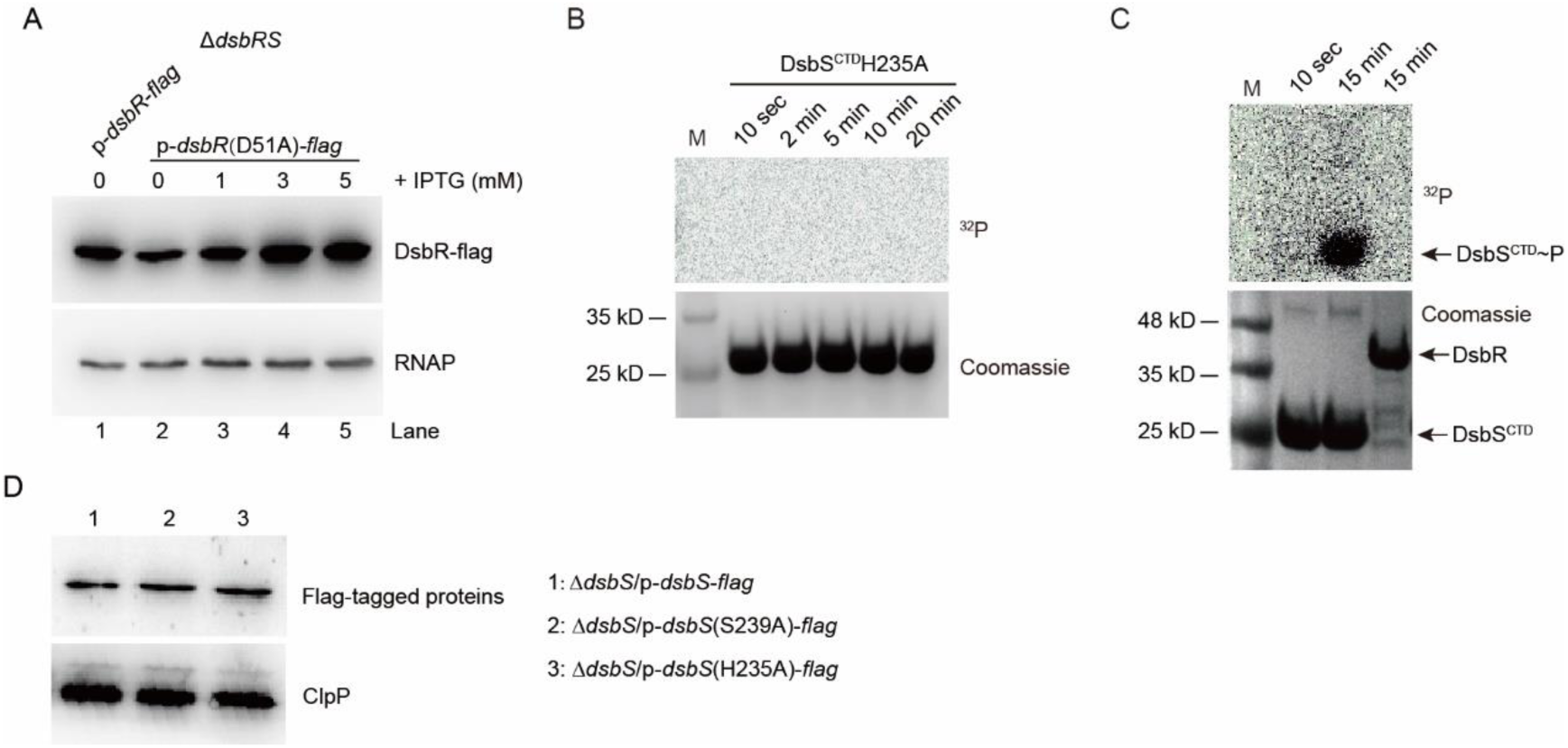
Western blot and phosphorylation assays. (A) Western blotting analysis for the production of DsbR-flag and DsbR(D51A)-flag in Δ*dsbRS* mutant grown in LB supplemented with indicated concentrations of IPTG at 37°C for 5 h. RNA polymerase (RNAP) was used as a loading control. p-*dsbR*-*flag* denotes pRK415-*dsbR*-*flag*, p-*dsbR*(D51A)-*flag* denotes pRK415-*dsbR*(D51A)-*flag*. (B) Time course of auto-phosphorylation of N-terminal His_6_-tagged DsbS^CTD^H235A mutant. M, protein marker. (C) The phosphorylation of SUMO-DsbR and DsbS^CTD^ incubated with [γ-^32^P]ATP for the indicated time. M, protein marker. (D) Western blotting analysis for the production of DsbS-flag, DsbS(H235A)-flag, and DsbS(S239A)-flag in Δ*dsbS* mutant grown in LB at 37°C for 12 h. ClpP was used as the loading control.

**Fig. S4.**
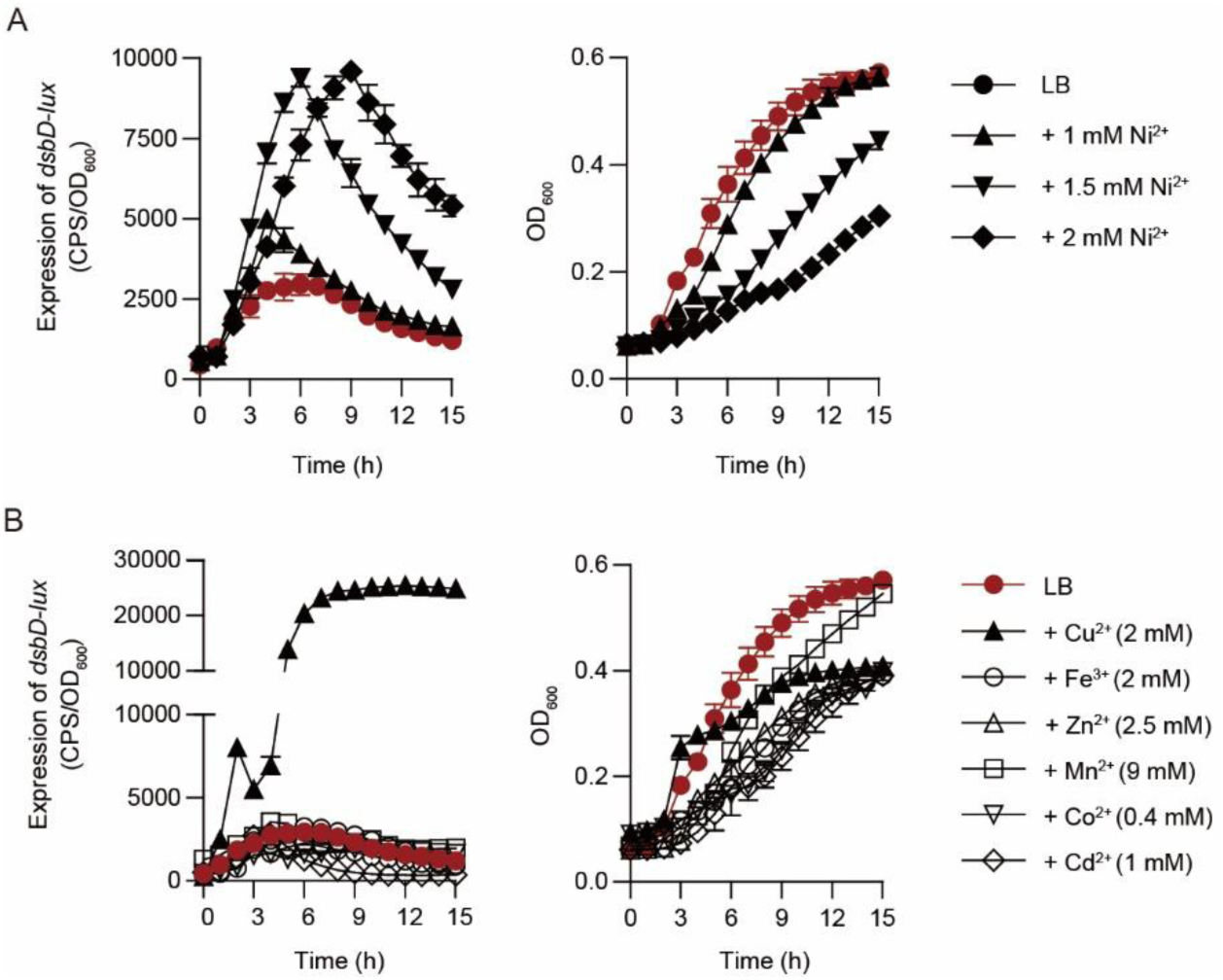
Determination of metal specificity of the *dsbD* promoter. (A) Expression of *dsbD*-*lux* (left panel) and the growth curve (right panel) of WT MPAO1 cultured in LB supplemented without or with indicated concentration of NiSO_4_. Data from n = 3 biological replicates reported as mean ± SD. (A) Expression of *dsbD*-*lux* (left panel) and the growth curve (right panel) of WT MPAO1 cultured in LB supplemented without or with indicated concentration of metal ions. The depicted metal salts and their concentrations are 2 mM CuSO_4_, 2 mM FeCl_3_, 2.5 mM ZnSO_4_, 9 mM MnSO_4_, 0.4 mM CoSO_4_, and 1 mM CdCl_2_. Data from n = 3 biological replicates reported as mean ± SD.

**Fig. S5.**
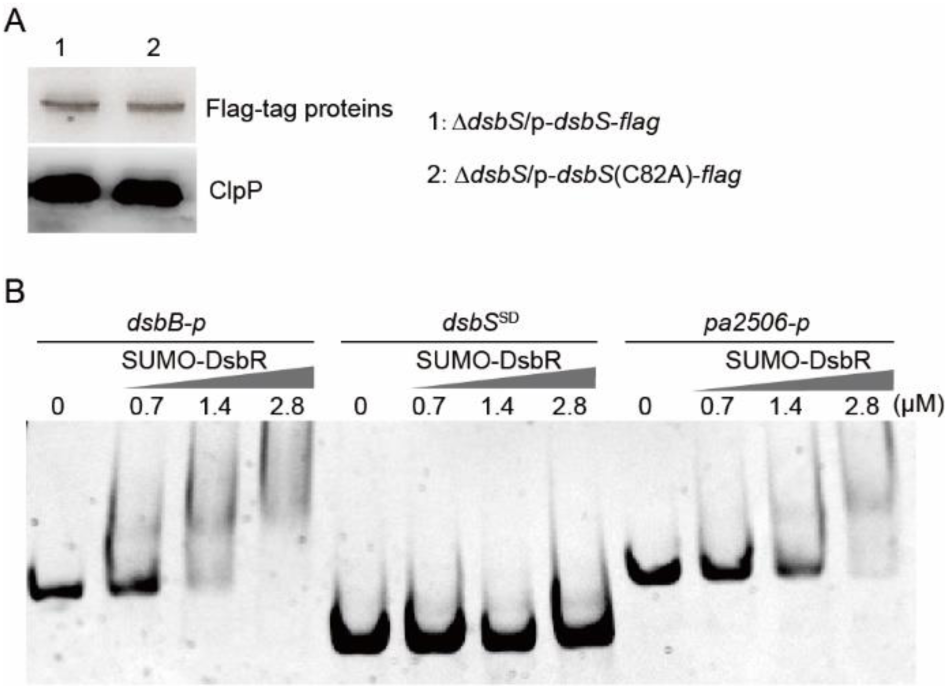
Western blot analysis and EMSAs. (A) Western blotting analysis for the production of DsbS-flag and mutant protein DsbS(C82A)-flag in Δ*dsbS* mutant grown in LB at 37°C for 12 h. ClpP was probed as a loading control. (B) EMSAs. DNA fragments, *dsbB*-*p*, *dsbS*^SD^, and *pa2506*-*p* were mixed with an increasing amount of purified SUMO-DsbR protein for EMSA assay, respectively, as indicated. *dsbB*-*p*, a 475-bp DNA fragment covering the promoter region of *dsbB*; *dsbS*^SD^, a 375-bp DNA fragment encoding the periplasmic domain region of DsbS; *pa2506*-*p*, a 478-bp DNA fragment covering the promoter region of *pa2506*.

**Fig. S6.**
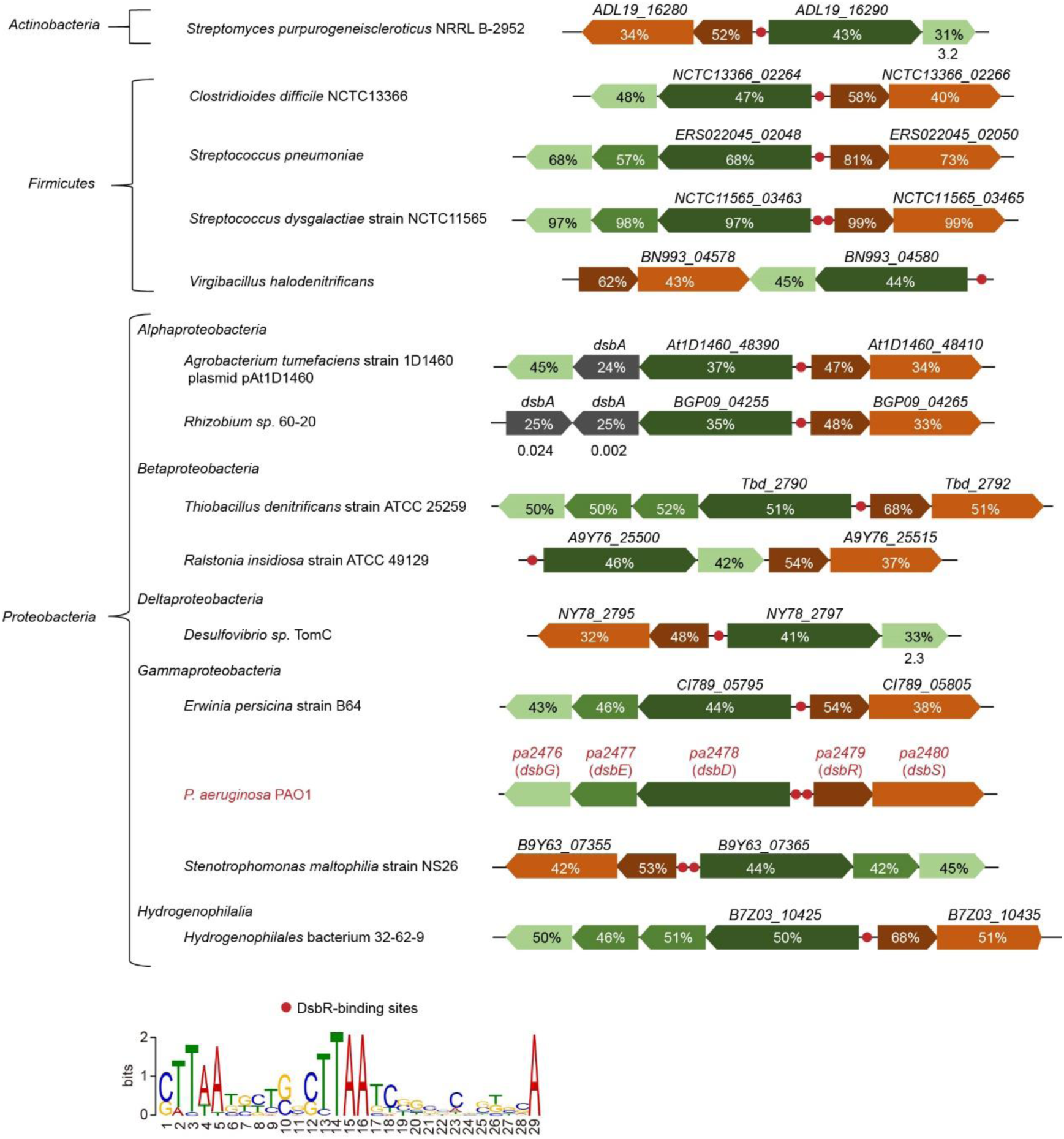
Genome context of *dsbRS* and its *dsbDEG* cognates in bacteria. A BLAST search revealed that the homologs of *P. aeruginosa* Dsb genes (i.e., *dsbD*, *dsbE*, and/or *dsbG*) are located adjacent to *dsbRS* in bacterial species belong to different phylum of bacteria including *actinobacteria*, *firmicutes*, and *proteobacteria*. Gene is shown by arrow and the candidate DsbR-binding site is shown by red circle. Genes with the same color represent homologous gene products. In all cases, Blastp E-value ≤ 10^−5^, except *dsbG* in *Streptomyces purpurogeneiscleroticus* NRRL B-2952 strain, *dsbA* in *Rhizobium sp.* 60-20 strain, and *dsbG* in *Desulfovibrio sp.* TomC strain, and the Blastp E-value of those gene products are indicated at the bottom of the arrows. The percentage of amino acid sequence identity to the corresponding proteins in *P. aeruginosa* MPAO1 are indicated in the arrows. The locus tags of the genes are indicated at the top of the arrows. For motif discovery, the 140 bp DNA sequences upstream of the start codon of DsbD homolog genes were subjected to MEME program with default settings except that the site distribution was set to any number of repetitions (anr).

**Fig. S7.**
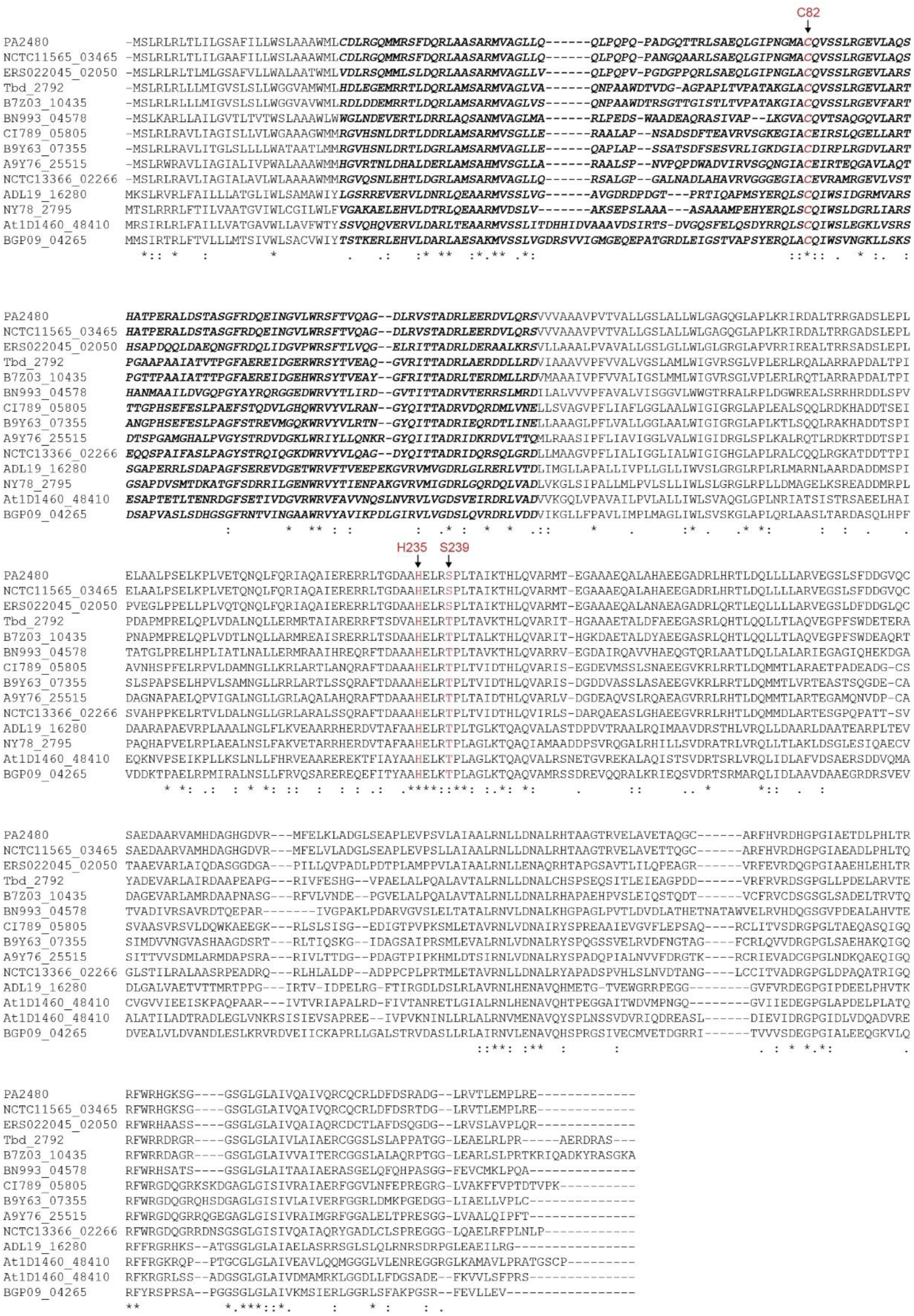
Sequence alignment of DsbS homologues. The sequences were aligned using the ClustalW program, and the locus tags of the DsbS homolog genes in different bacterial strains correspond to those described in Fig. S6. Stars indicate identical amino acids, double dots (:) indicate conserved amino acids, and single dots (.) indicate that residues are more or less similar. The periplasmic sensor domain of DsbS homologues (residues 28-146) is in bold and in italics. The three key residues (i.e., Cys82, His235, and Ser239) inferred from this study are highlighted in red.

**Fig. S8.**
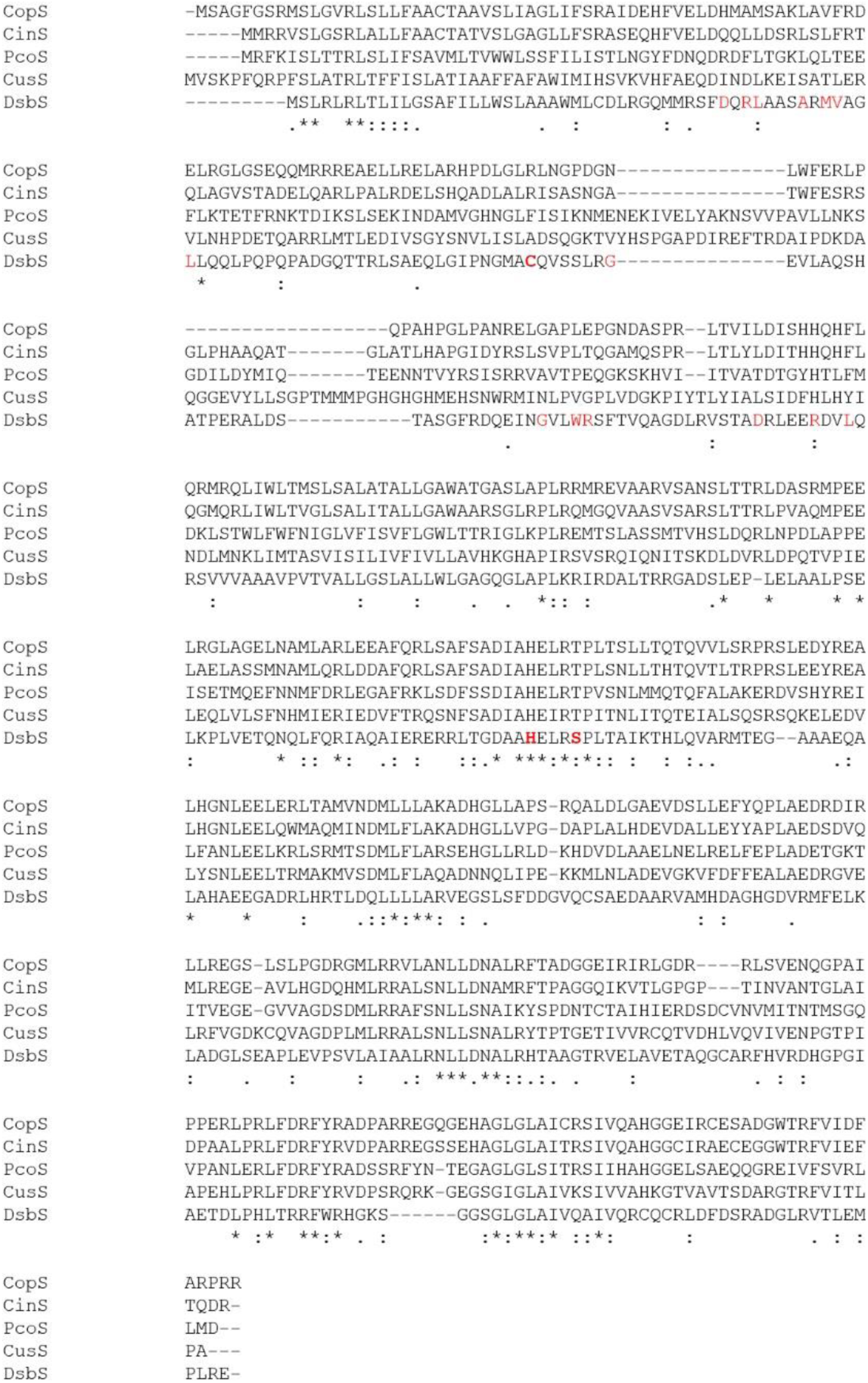
Sequence alignment of DsbS and copper-sensing HKs CopS, CinS, PcoS and CusS. Asterisks indicate residues exactly conserved across all the sequences, while dots indicate similar amino acids. The Cys82, His235, and Ser239 residues of *P. aeruginosa* DsbS are depicted in bold red. The identical amino acids in the periplasmic domain of DsbS homologs (Fig. S8) are shown in red. CopS, the PA2810 of *P. aeruginosa* PAO1; CinS, the PP_2157 of *Pseudomonas putida* KT2440; PcoS, the PcoS of *Klebsiella pneumoniae* TUM18984 (plasmid pTHC11-1); CusS, the PRK09835 of *Escherichia coli* str. K-12 *substr*. MG1655.

### Tables

**Table S1.**
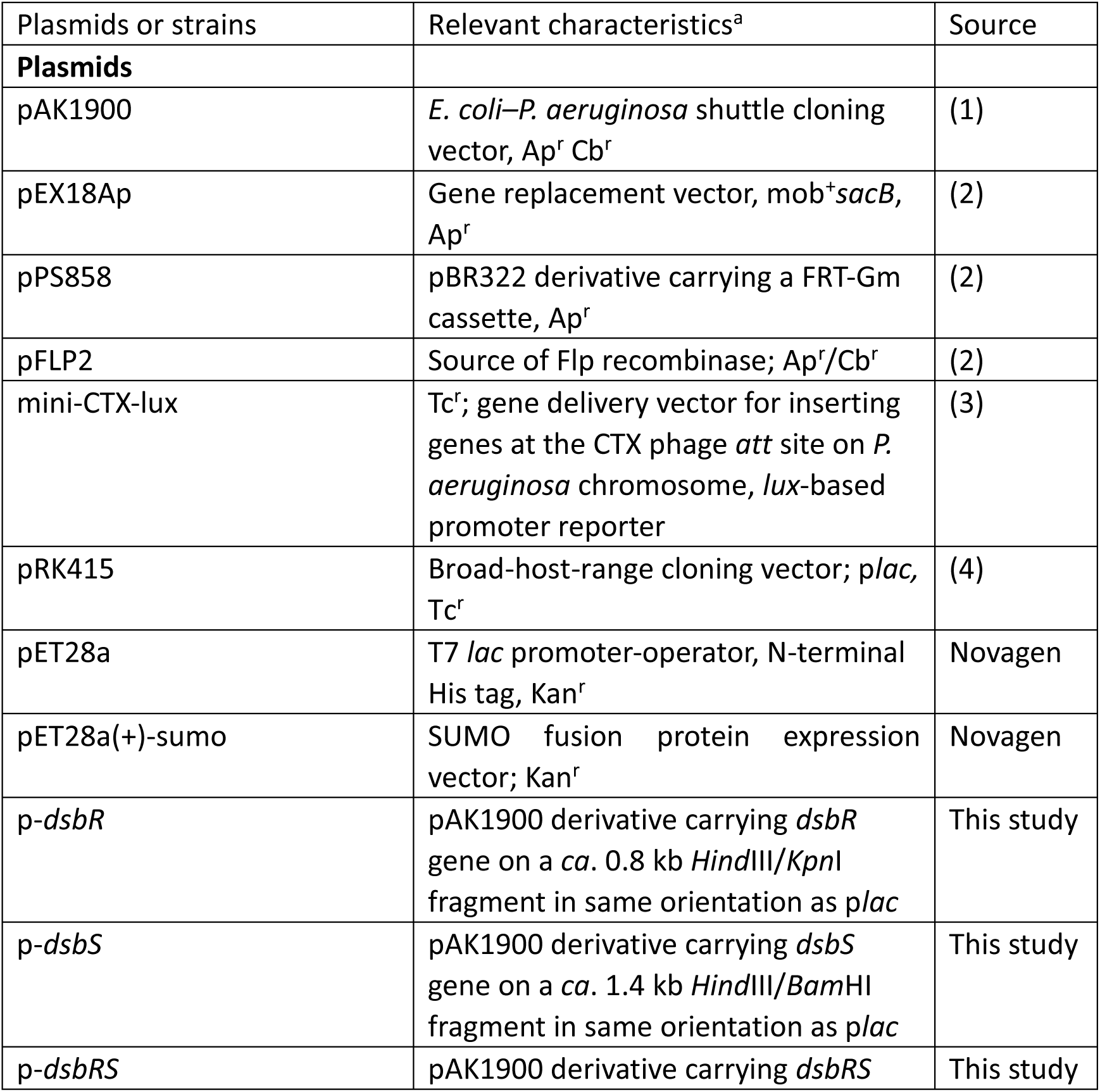
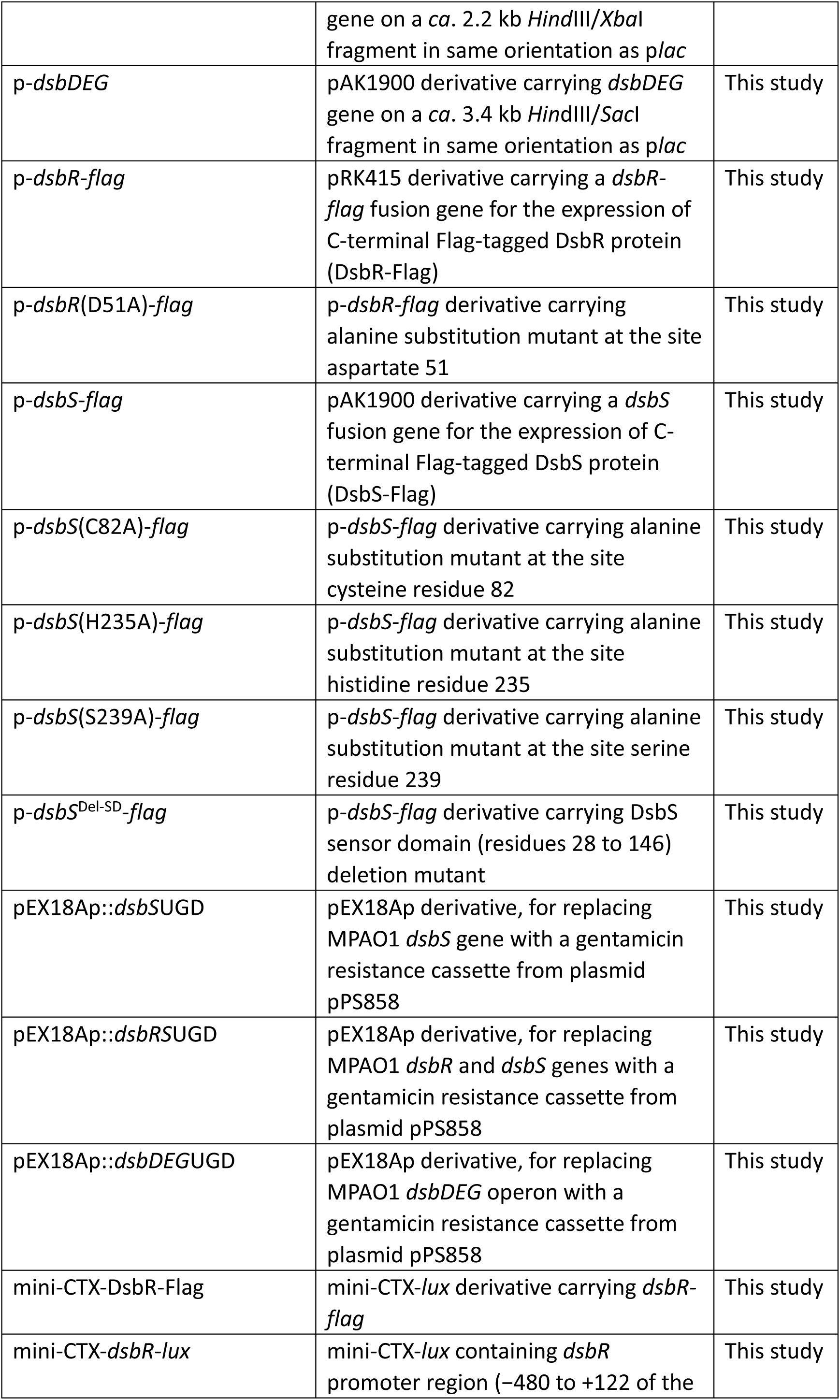
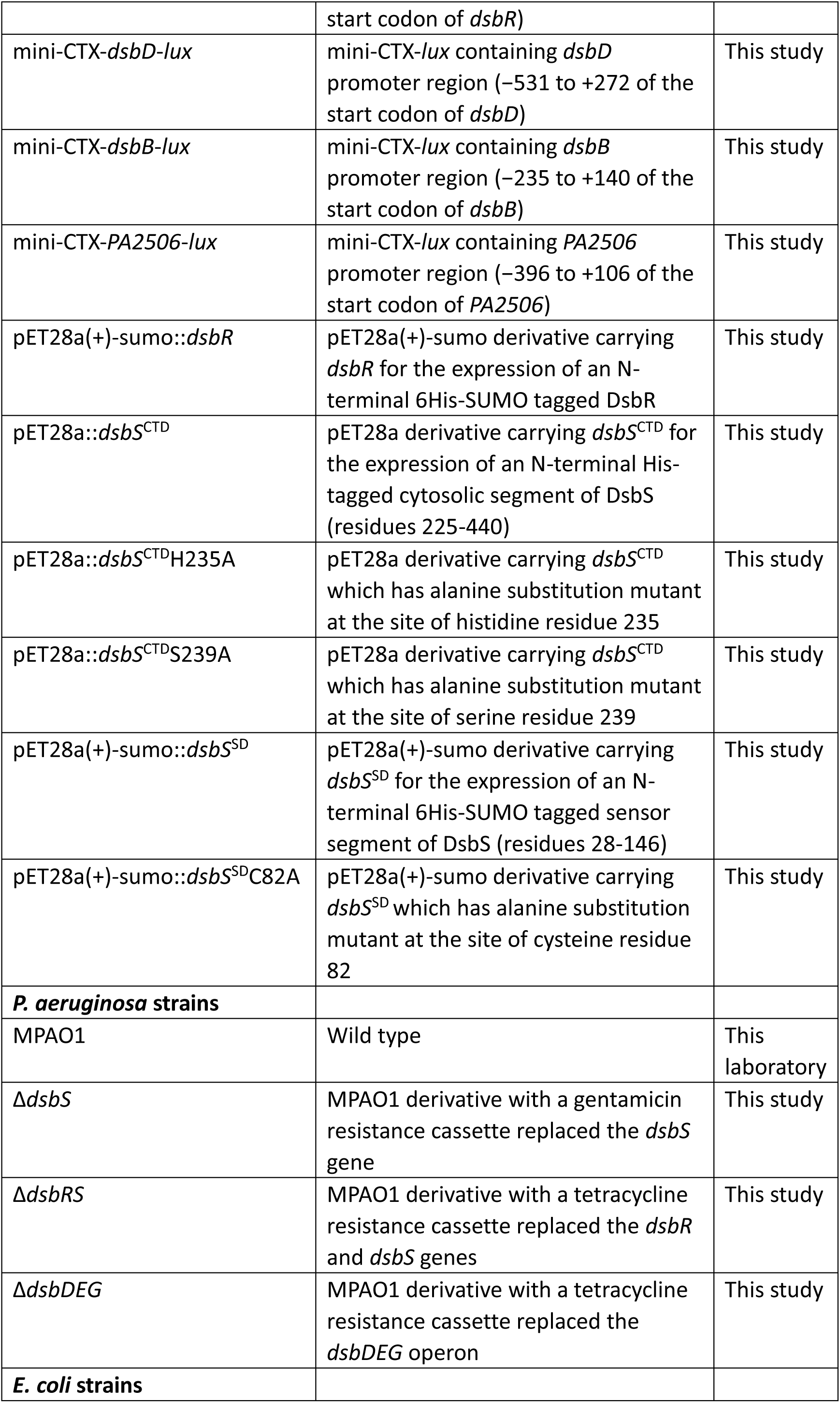
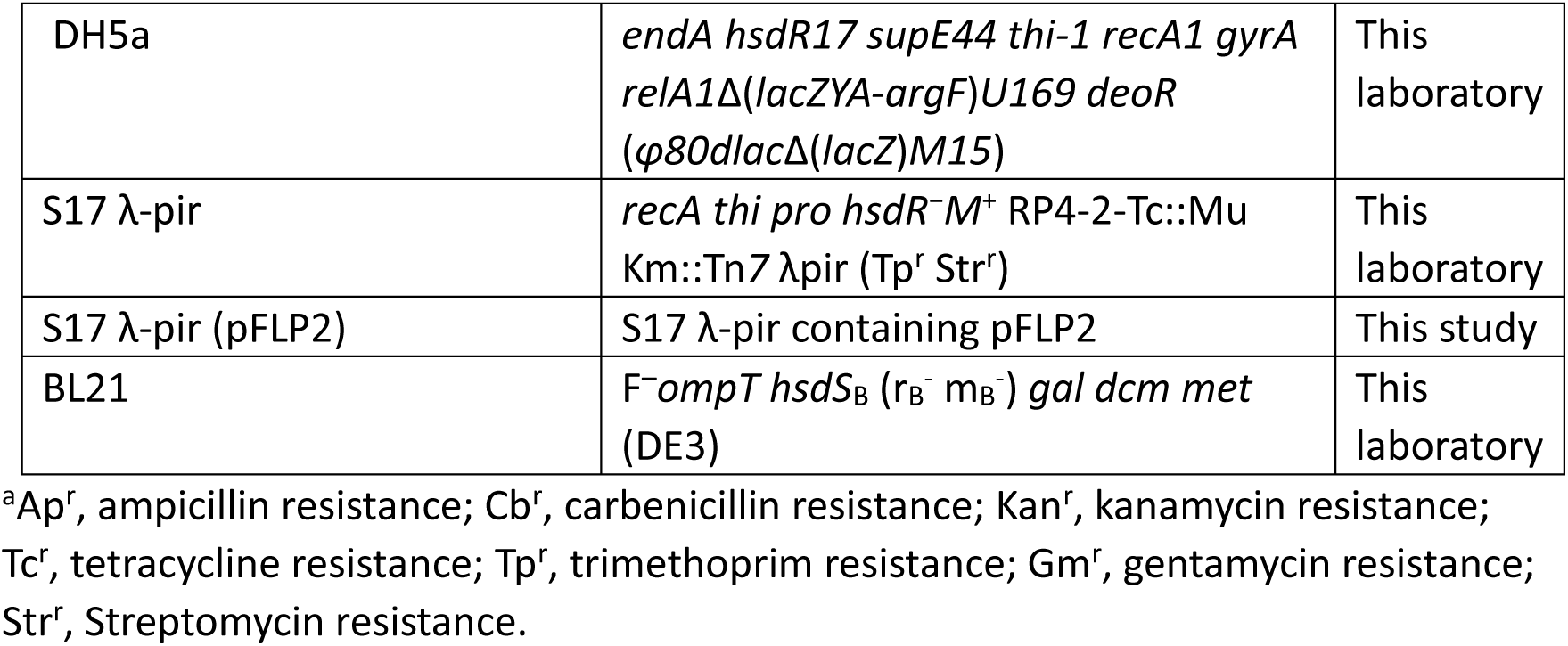
Plasmids and bacterial strains and used in this study

**Table S2.**
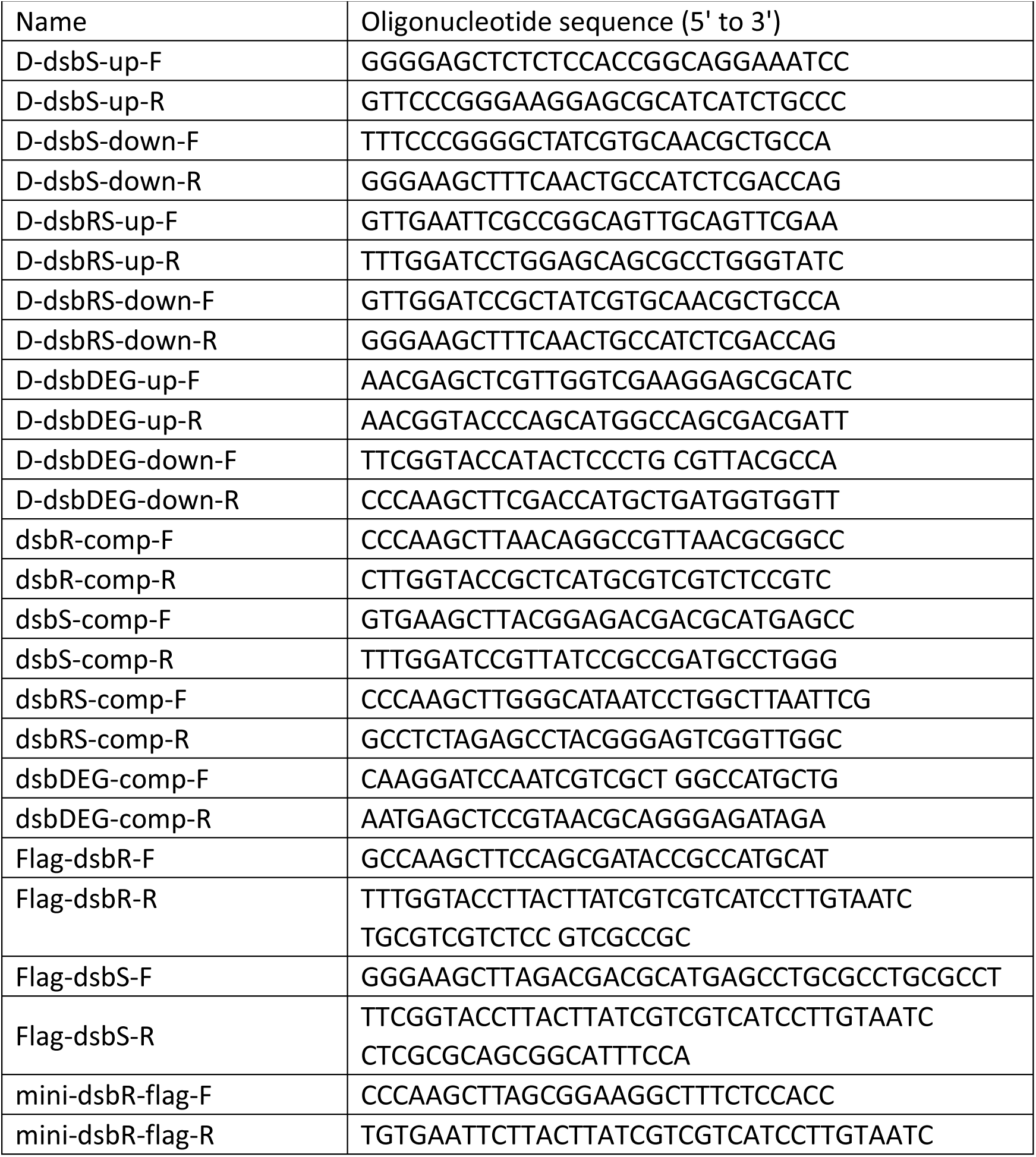
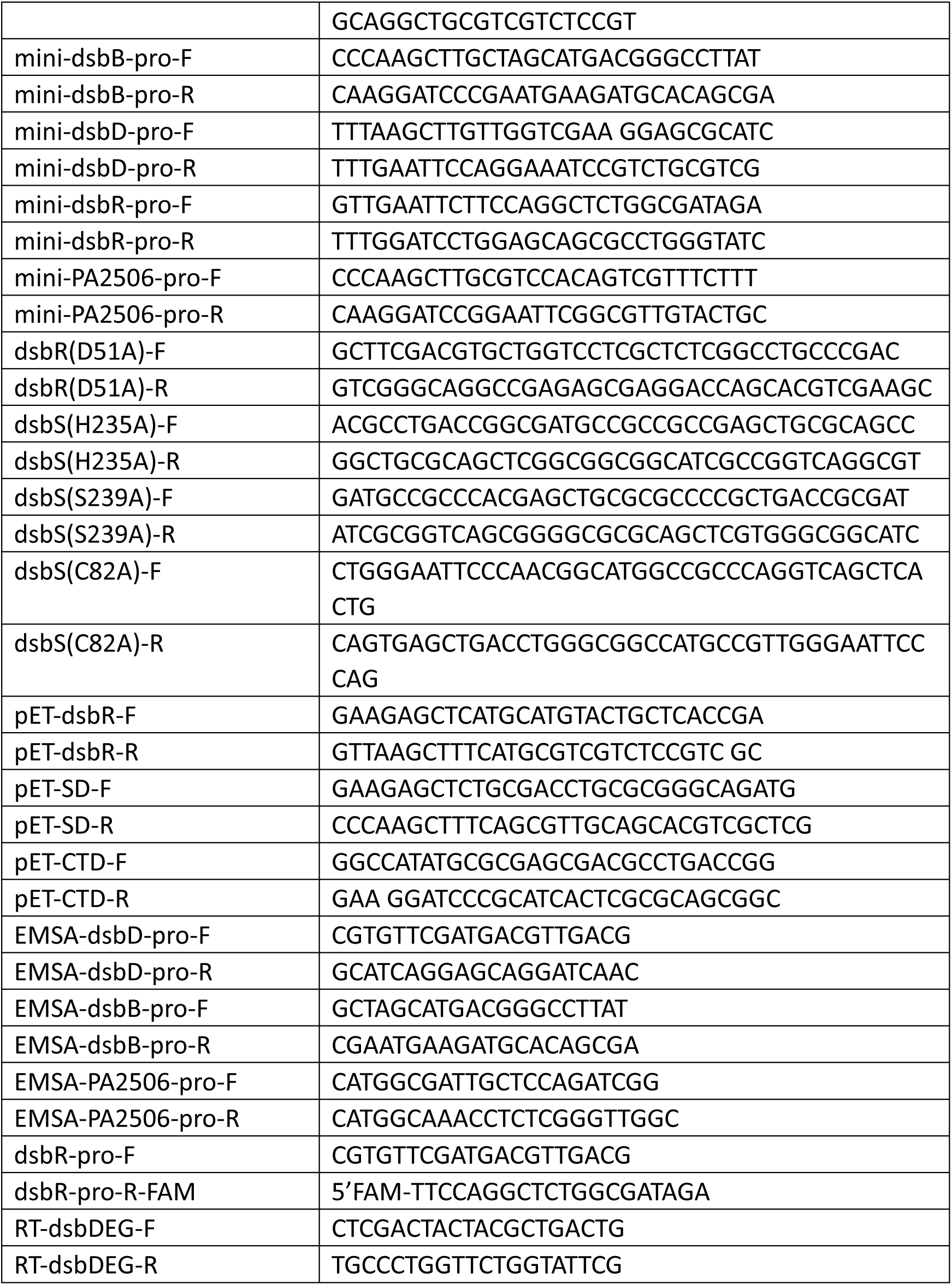
Primers used in this study

